# Clinical and pathological characterization of Central Nervous System cryptococcosis in an experimental mouse model of stereotaxic intracerebral infection

**DOI:** 10.1101/2022.10.31.514648

**Authors:** Mohamed F. Hamed, Vanessa Enriquez, Melissa E. Munzen, Claudia L. Charles-Niño, Mircea R. Mihu, Habibeh Khoshbouei, Karina Alviña, Luis R. Martinez

**Author notes:** To whom correspondence should be addressed: Luis R. Martinez, University of Florida College of Dentistry, Department of Oral Biology, 1395 Center Drive, DG-48, P.O. Box 100424, Gainesville, FL 32610.

## Abstract

Infection of the Central Nervous System (CNS) by the encapsulated fungus *Cryptococcus neoformans* can lead to high mortality meningitis, most commonly in immunocompromised patients. While the mechanisms by which the fungus crosses the blood-brain barrier to initiate infection in the CNS are well recognized, there are still substantial unanswered questions about the disease progression once the fungus is established in the brain. *C. neoformans* is characterized by a glucuronoxylomannan (GXM)-rich polysaccharide capsule which has been implicated in immune evasion, but its role during the host CNS infection needs further elucidation. Therefore, the present study aims to examine these key questions about the mechanisms underlying cryptococcal meningitis progression and the impact of fungal GXM release by using an intracerebral rodent infection model via stereotaxic surgery. After developing brain infection, we analyzed distinct brain regions and found that while fungal load and brain weight were comparable one-week post-infection, there were region-specific histopathological (with and without brain parenchyma involvement) and disease manifestations. Moreover, we also observed a region-specific correlation between GXM accumulation and glial cell recruitment. Further, mortality was associated with the presence of subarachnoid hemorrhaging and GXM deposition in the meningeal blood vessels and meninges in all regions infected. Our results show that using the present infection model can facilitate clinical and neuropathological observations during the progression of neurocryptococcosis. Importantly, this mouse model can be used to further investigate disease progression as it develops in humans.

**IMPORTANCE:** *Cryptococcus neoformans* is the causative agent of cryptococcal meningoencephalitis (CME), a severe infection of the central nervous system (CNS) in humans. The high mortality and morbidity rates of this disease in immunosuppressed and HIV^+^/AIDS-infected individuals necessitate a reliable model for understanding the mechanisms and manifestations of cryptococcosis infection in the CNS. This study presents a mouse model of infection that utilizes a stereotaxic surgical setup to provide a comparative analysis of *C. neoformans* infection in the brain. Though not typical of infection in humans, this method provides a controlled experimental system to observe the progression of *C. neoformans* infection in specific regions of the brain. Additionally, this study characterizes murine manifestations of the disease, including GXM deposition in brain tissue and blood vessels, and documents the role of microglia, the resident immune cells of the CNS, in engulfing fungal cells. Depending on the region of the brain infected, distinct disease pathology can be observed with this model, expanding our understanding of cryptococcosis progression in the CNS. The importance of this work extends to the clinical setting, where many questions regarding cerebral cryptococcosis remain unresolved. Utilization of this model to further understand the mechanisms of this pathogen will provide means to develop new therapies to combat infection in humans.

## INTRODUCTION

*Cryptococcus neoformans* is an encapsulated opportunistic yeast-like fungus that triggers life-threatening meningoencephalitis in both immunocompromised and healthy individuals. Of the ~220,000 annual cases of cryptococcal meningitis in HIV^+^/AIDS victims worldwide, 181,000 (~82%) resulted in death (1). *Cryptococcus* species are the third most common invasive fungus in solid organ transplantation (SOT) recipients (2). The prevalence of cryptococcosis in this population ranges between 0.2 to 5.8%, with a total mortality rate of 20% to 50%, highlighting its prevalence in another patient population at risk (2, 3). *C. neoformans* infects humans when desiccated or poorly encapsulated yeasts/basidiospores are inhaled (4, 5). Most immunocompetent hosts render *C. neoformans* dormant in their lungs, but immunosuppression can trigger its replication and dissemination via blood or lymph to other organs, especially the brain. Approximately 50%-75% of SOT recipients with cryptococcosis have extrapulmonary or disseminated disease (6, 7). During cryptococcosis, fungemia is detected in ~50% of HIV-infected patients (8), correlates experimentally with dissemination and brain invasion (9), and is an independent parameter of early mycological failure in humans (8). *C. neoformans* crosses the blood-brain barrier (10) via multiple mechanisms: transcytosis, paracellular transit, or as “Trojan horse” cargo within host inflammatory cells (11–14). Once in the brain, *C. neoformans* triggers meningoencephalitis, the most serious hallmark of cryptococcosis.

With a protective polysaccharide capsule, *C. neoformans* grows at mammalian physiological temperature and produces melanin to aid in virulence (15). The capsule is the most distinctive physical structure of the cryptococcal cell and contributes substantially to its virulence (16–18). Compared to *in vitro* conditions, most *C. neoformans* strains form larger capsules in tissue (17, 19), where capsule size can be influenced by physiological CO_2_ levels (20), serum (21), osmolarity (22, 23), and iron levels (24). Glucuronoxylomannan (GXM) is the primary constituent of the capsule and accumulates in the serum and cerebrospinal fluid (CSF) (25, 26). High GXM levels are associated with many immunosuppressive effects, including interference with phagocytosis, antigen presentation, leukocyte migration and proliferation, and specific antibody (Ab) responses, contributing directly to *C. neoformans* pathogenesis (27). GXM even enhances HIV replication (28), but despite this knowledge, many specific effects of GXM on *C. neoformans* brain pathogenesis or inflammatory response remain largely unresolved.

Cryptococcal meningoencephalitis (CME) is uniformly fatal if left untreated, and even with optimal antifungal treatment, mortality ranges from 30–100% (29–31). Non-HIV CME patients display greater brain tissue inflammation and fewer extracellular cryptococci than HIV-associated CME victims (32). The latter group displays numerous cryptococci in the subarachnoid space post-mortem, associated with raised intracranial pressure (ICP) and hydrocephalus (33), CSF fungal burden, and GXM accumulation, which are all associated with mortality (34). In addition, blood vessel occlusion and damage are frequently associated with systemic infection, given that individuals with CME develop aneurisms or subarachnoid hemorrhage (35). In fact, there are instances where CME has been associated with ischemic stroke with vasculitis in HIV^+^ (36) and HIV^-^ individuals (37). Nevertheless, the mechanisms of CME pathogenesis and the progression of its manifestations are poorly understood.

In this study, we characterized the clinical and pathological manifestations of CNS cryptococcosis in an experimental mouse model of intracerebral (i.c.) infection using stereotaxic surgery. Our results showed that depending on the brain region where *C. neoformans* infection was found, cerebral cryptococcosis develops with or without brain parenchyma involvement. *C. neoformans*-infected mouse brains displayed marked similarities with histopathological and clinical manifestations observed in humans, making our model an excellent method to distinguish the direct neurotoxic impact of fungal infection, GXM release, and disease progression. Future research using this mouse model of i.c. fungal infection may help fill in the knowledge gap in our understanding of how cerebral cryptococcosis progresses and may provide novel research avenues in *C. neoformans* pathogenesis.

## RESULTS

### i.c. *C. neoformans* infection in different brain regions of C57BL/6 mice results in hydrocephalus, subarachnoid hemorrhage, and death

We investigated the development of cerebral cryptococcosis after i.c. *C. neoformans* infection (10^4^ fungi of strain H99 in 1 μL of sterile saline) in C57BL/6 mice. We injected the fungus in different regions of the brain including the midline at bregma level, neocortex, striatum, and ventricles (*n*=10 mice per region, Fig. 1A) using a stereotaxic apparatus (Supplemental Fig. 1). Bregma is an anatomical landmark in the skull where the sagittal and coronal sutures intersect joining the parietal and frontal bones [coordinates of injection: medial/lateral (X-axis): 0, anterior/posterior (Y-axis): 0, dorsal/ventral (Z-axis): −2.5); Table 1; Fig. 1A, upper left panel]. The cerebral cortex is the largest site of neural integration in the CNS playing key roles in attention, perception, awareness, memory, language, and consciousness [coordinates of injection: medial/lateral (X-axis): +/-1.5, anterior/posterior (Y-axis): 0, dorsal/ventral (Z-axis): −1); Table 1; Fig. 1A, upper right panel]. The striatum serves as the primary input to the rest of the basal ganglia, receives glutamatergic and dopaminergic inputs from different sources, and is a critical component of the motor and reward systems [coordinates of injection: medial/lateral (X-axis): +/-2, anterior/posterior (Y-axis): 0.2, dorsal/ventral (Z-axis): −3.5); Table 1; Fig. 1A, lower left panel]. The lateral ventricles of the brain are a communicating network of cavities filled with CSF and located within the brain parenchyma [coordinates of injection: medial/lateral (X-axis): +/-1.1, anterior/posterior (Y-axis): 0.5, dorsal/ventral (Z-axis): −2.5; Table 1; Fig. 1A, lower right panel]. Mice infected in the cerebral cortex and right ventricle showed similar and the earliest mortality [median survival, 8-dpi; *P* < 0.05] (Fig. 1B). The clinical symptoms shown by mice in these groups include respiratory distress, dehydration, severe hunching, dyspnea with considerable abdominal involvement, abnormal neurologic movement with consistent body twitching, and body condition score of 2.5 (Table 2). All the animals infected in bregma died by 12-dpi. These mice experienced dehydration, hunching, lethargy, body condition score of 2, and a firm cranial deformity of 1.3 mm in diameter (Table 2). Mice infected in the striatum survived the longest, 20-dpi with a median survival of 15-dpi (Fig. 1B). Mice infected in the basal ganglia demonstrated a plethora of clinical signs likely due to their prolonged survival including respiratory distress, mild dehydration, severe hunching, dyspnea, palpitations, locomotor disturbance and rapid movement when alert, recumbency with spastic limb movement (Table 2). Mice infected with *C. neoformans* in specific brain regions (*n* = 5 per group) exhibited similar fungal burden 3- (bregma, 3.38 × 10^2^ colony forming units (CFU) per gram (g)/tissue; cortex, 3.92 × 10^2^ g/tissue; striatum, 4.47 × 10^3^ g/tissue; ventricle, 1.43 × 10^3^ g/tissue) and 7- (bregma, 3.31 × 10^4^ CFU per g/tissue; cortex, 2.39 × 10^4^ g/tissue; striatum, 4.29 × 10^4^ g/tissue; ventricle, 5.7 × 10^4^ g/tissue) dpi, regardless of the time of death of all the animals within a specific group (Fig. 1C). C57BL/6 rodents infected with *C. neoformans* in the striatum developed signs of hydrocephalus at 12 to 14 dpi such as bulging of the forehead (black arrows; right panel) and the dome-shaped cranium relative to sterile saline injected and uninfected control animals (saline injected/uninfected control; left panel; Table 2; Fig. 1D). A representative brain coronal tissue section of saline injected/uninfected mice displayed normal lateral ventricles (black arrows), septum pellucidum (black arrowhead), and parenchyma (Fig. 1E). In contrast, brain tissue from infected groups with *C. neoformans* evinced obvious signs of hydrocephalus including ventriculomegaly in both lateral and third (red arrows) ventricles, thinning of the cerebral cortex (dashed circles), and widely distributed pale and red microinfarcts throughout the brain parenchyma. Mice infected in bregma, cortex, and striatum also showed cryptococcoma or *C. neoformans* brain-lesion formation and expansion in tissue due to encephalomalacia or liquefactive necrosis. Gross morphology of representative brains excised from mice infected in bregma (B), cortex (C), striatum (S), and right ventricle (V) upon termination (day 7-dpi for B, C, and V and 20-dpi for S) after perfusion and fixation with 4% paraformaldehyde revealed swollen cerebral hemispheres with compressed olfactory bulb, subarachnoid hemorrhage, clotted blood at the superior sagittal sinus and transverse sinus (Fig. 1F), in comparison to saline injected/uninfected control (Ctr) mice (Fig. 1F). Periodic acid-schiff (PAS)-stained sections of the excised brains 7-dpi were performed and confirmed the presence of cryptococci (black arrowheads, red staining) in the brain-lesions (Fig. 1G). Finally, weight determinations of brains removed 7-dpi from mice infected with *C. neoformans* in bregma (*P* < 0.01), cortex (*P* < 0.001), striatum (*P* < 0.001), or right ventricle (*P* < 0.0001) demonstrated a significant increase in tissue mass relative to saline injected/uninfected control mice. There were no differences in brain weight among the infected groups. Our data demonstrate that mice directly infected with *C. neoformans* in specific brain regions develop signs of hydrocephalus and subarachnoid hemorrhage resulting in mortality.

**Fig. 1.**
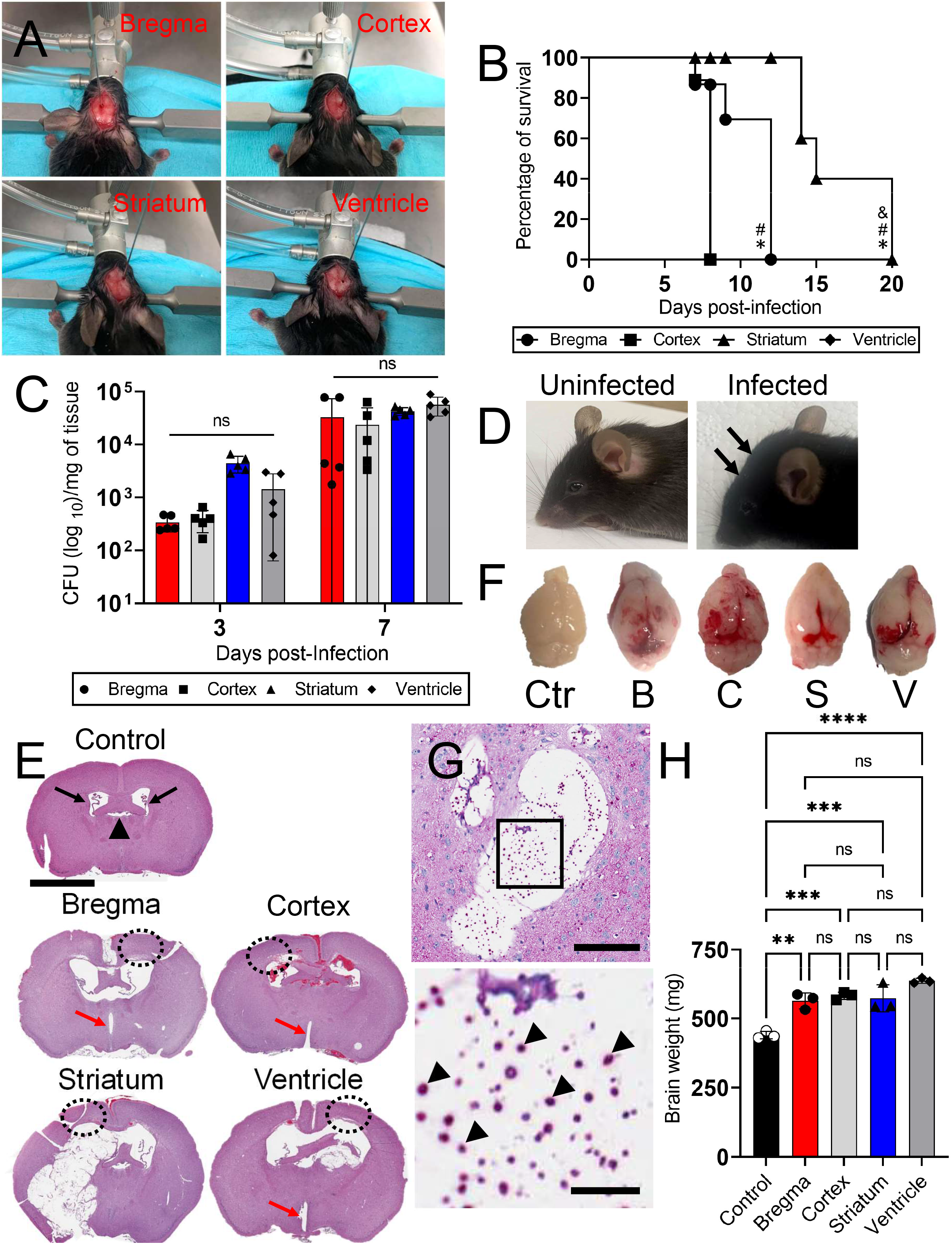
Intracerebral (i.c.) *Cryptococcus neoformans* infection of C57BL/6 mice in specific brain regions causes hydrocephalus and subarachnoid hemorrhage exacerbating mortality. **(A)** C57BL/6 mice of 6-8 weeks old were i.c. infected with a single inoculum of 10^4^ *C. neoformans* strain H99 cells in either bregma, cortex, striatum (basal ganglia), or ventricle after a stereotaxic surgery. **(B)** Survival differences of C57BL/6 mice infected with 10^4^ fungi (*n*=10 mice per group) in bregma (circle), cortex (square), striatum (triangle), or ventricle (diamond). Symbols (*, #, and &) indicate *P* value significance (*P* < 0.05) calculated by log rank (Mantel-Cox) analysis. *, #, and & indicate significantly longer survival than in the cortex-, ventricle-, and bregma-infected groups, respectively. **(C)** Fungal burden in whole brain tissue (numbers of colony forming units (CFU)/milligram (mg) of tissue) collected from bregma-, cortex-, striatum-, and ventricle-infected mice with 10^4^ cryptococci (*n* = 5 per group) 3- and 7-days post-infection (dpi). **(D)** Photograph of a mouse infected with *C. neoformans* H99 in the striatum of the basal ganglia showing signs of hydrocephalus (right) including bulging of the forehead (black arrows). A picture of a healthy uninfected mouse (left) is shown for comparison. These animals were euthanized 14-dpi. **(E)** Representative brain coronal histology sections of uninfected control and bregma-, cortex-, striatum-, and ventricle-infected mice. Uninfected mice exhibit normal brain parenchyma, regular lateral ventricles (black arrows) and an intact septum pellucidum (arrowhead). Dashed black ellipses demonstrate thinning of the cortex. Red arrows indicate ventriculomegaly of the third ventricle. Scale bar: 2 mm. **(F)** Gross morphology of control and bregma-, cortex-, striatum-, and ventricle-infected brains after perfusion with 4% paraformaldehyde. For **E** and **D**, uninfected (control) and infected mice in bregma, cortex, and ventricle were euthanized 7-dpi. Striatum-infected rodents were sacrificed 20-dpi. **(G)** Representative periodic acid-schiff (PAS; fungi; red staining; scale bar: 100 μm)-stained sections of the brain are shown and were removed 7-dpi. Lower panel image is a magnification of the smaller box in the upper PAS-stained section to better show the cryptococcal cells (indicated with arrow heads; scale bar: 20 μm). **(H)** Brain weight determinations from tissue removed 7-dpi. For **C** and **H**, bars and error bars denote the means and standard deviations (SDs), respectively. Each symbol represents values for an individual mouse (*n* = 5 for **C** and *n* =3 for **H**). Asterisks denote *P*-value significance (\*\**P* <0.01, \*\*\**P* < 0.001, and \*\*\*\**P* < 0.0001) calculated using analysis of variance (ANOVA) and adjusted by use of the Tukey’s post-hoc analysis. ns denotes not statistically significant comparisons. These experiments were performed twice, similar results were obtained each time, and all the results combined are presented.

**Table 1.**
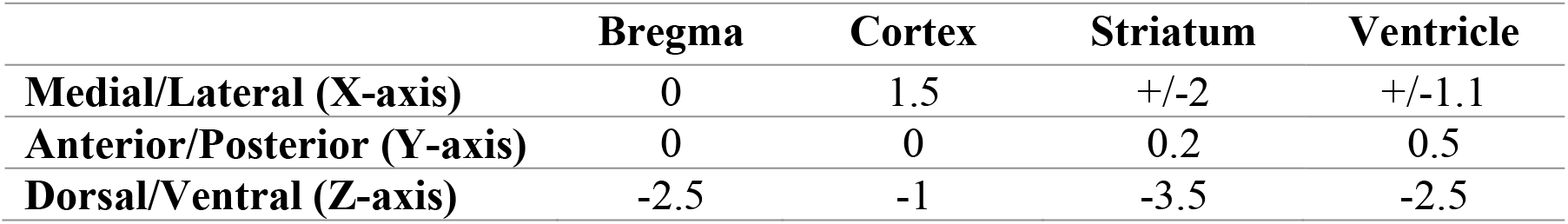
Stereotaxic coordinates for *C. neoformans* intracerebral infection of C57BL/6 mice.

**Table 2.**
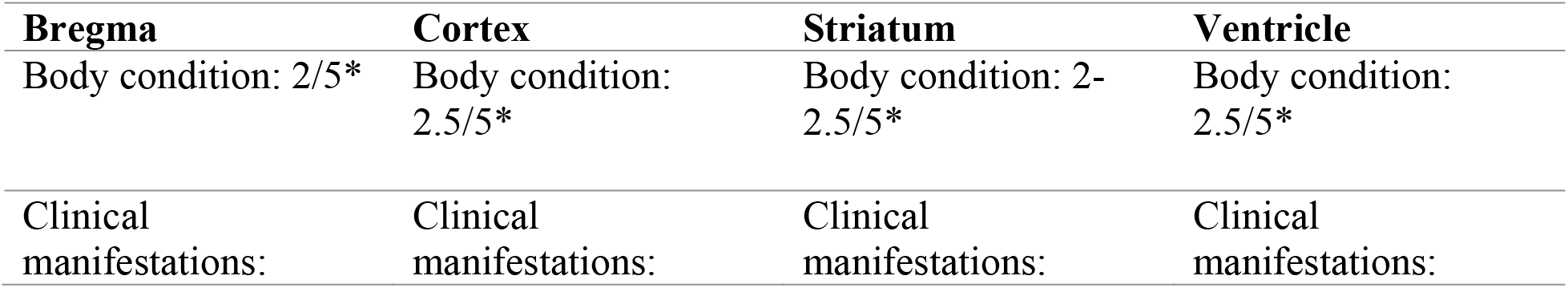
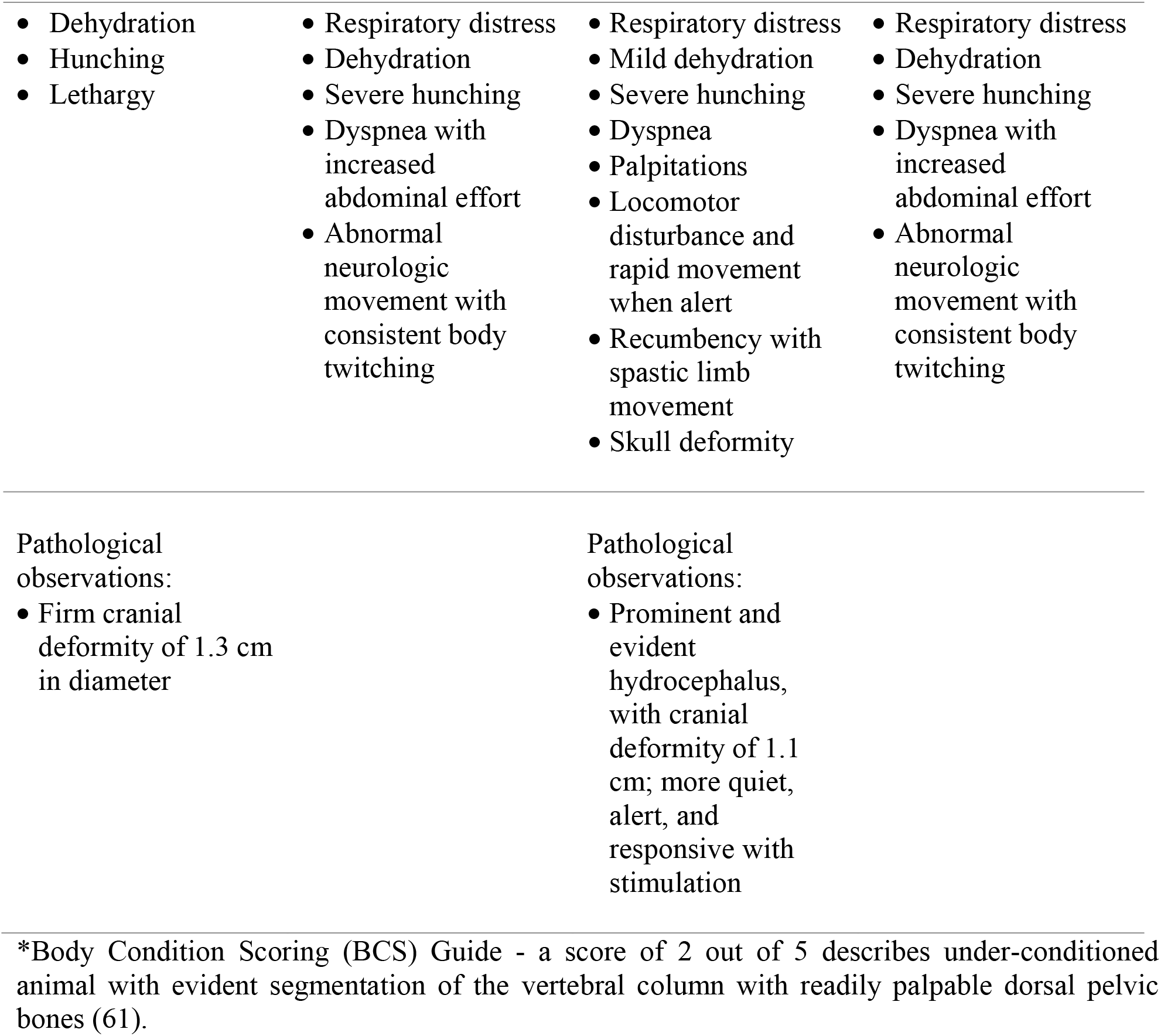
Clinical manifestations in mice with cerebral cryptococcosis after infection in specific brain regions.

### Characterization of cerebral cryptococcosis in mice infected in specific brain regions

We described the development of cerebral cryptococcosis in coronal brain sections excised from C57BL/6 mice directly infected in bregma 7-dpi. Mice exhibited encapsulated yeast cells lodged in the septum pellucidum inducing encephalomalacia and scattered microinfarcts in superficial (e.g., cerebral cortex and corpus callosum) and deep tissue (e.g., thalamus, hypothalamus, and basal ganglia) (Fig. 2A). Higher magnification images showed expanding encephalomalacia or liquefactive necrosis in the brain parenchyma (Fig. 2B) and minimal inflammation (Fig. 2C). GXM-stained immunohistochemistry (IHC) using monoclonal antibody (mAb) 18B7 (Fig. 2D), which is specific to the main component of *C. neoformans* capsular polysaccharide (CPS), revealed wide distribution of GXM especially in the cortical region above of each ventricle, septum pellucidum (encephalomalacia; Fig. 2E-F), and adjacent blood vessels, particularly in the margins of microinfarcts.

**Fig. 2.**
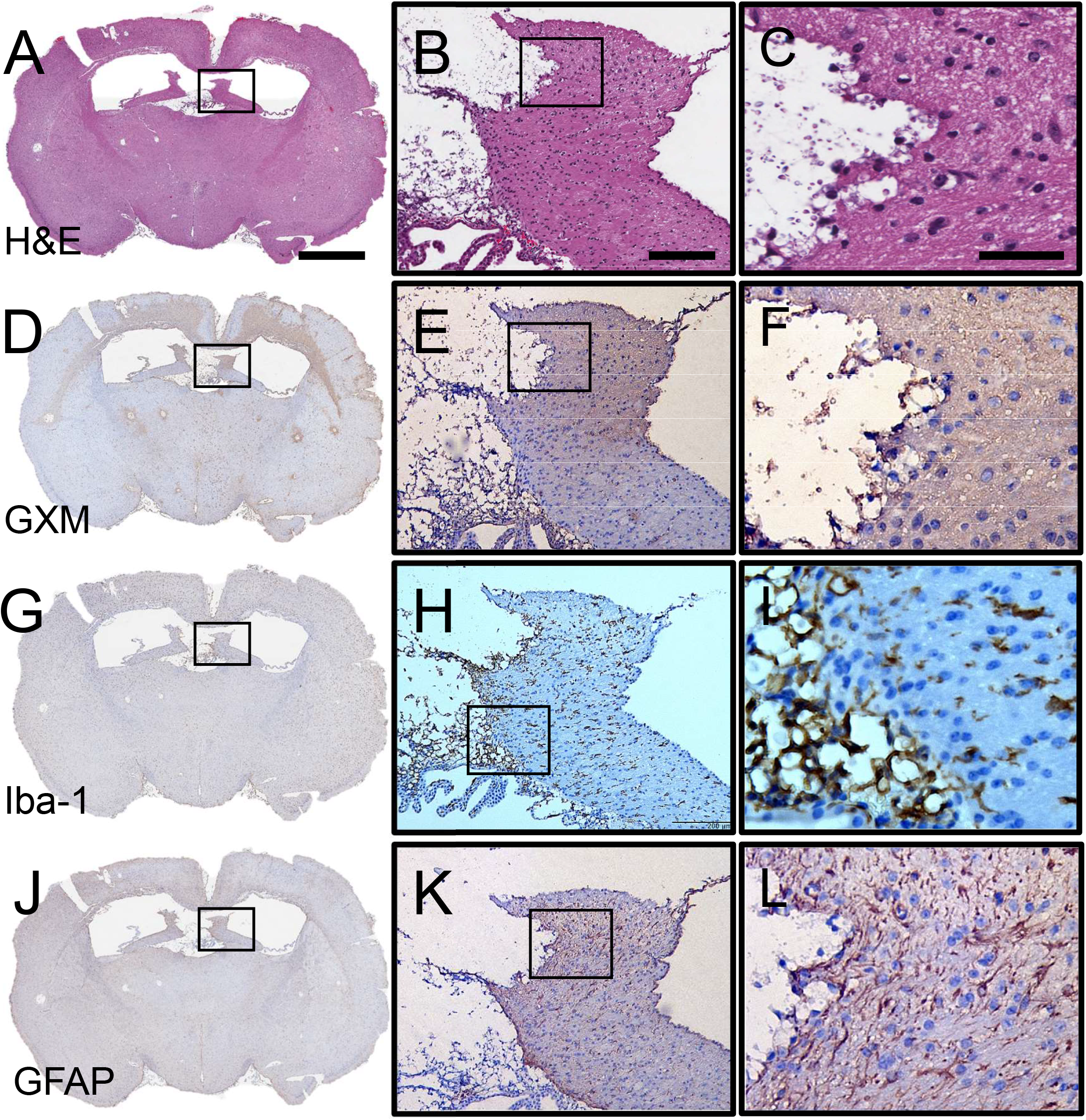
Cerebral cryptococcosis in mice infected in bregma. Representative coronal brain sections (scale bar: 1 mm) of bregma-infected mice stained with **(A-C)** hematoxylin and eosin (H&E), **(D-F)** *C. neoformans* glucuronoxylomannan (GXM)-specific monoclonal antibody (mAb) 18B7, **(G-I)** ionized calcium binding adaptor molecule 1 (Iba-1; microglia), and **(J-L)** glial fibrillary acidic protein (GFAP; astrocytes) are shown. Mouse brains were excised 7-dpi. Middle (scale bar: 100 μm) and right (scale bar: 20 μm) panel images are a magnification of the smaller box in the corresponding left-stained section to better show the tissue morphology, GXM distribution (brown stain), and glial cell recruitment/activation (brown stain). Five mice per infection route were evaluated to confirm all the histopathological observations reported in these studies.

We also examined glial responses (microglia and astrocytes) to the fungus at 7-dpi. IHC against the microglia-specific marker ionized calcium binding adaptor molecule 1 (Iba-1, Fig. 2G-I), demonstrated microglial response against the progressive encephalomalacia and cryptococcal growth (Fig. 2H). Microglia near fungal cells exhibited a dystrophic or necrotic morphology (Fig. 2L). Similarly, IHC against astrocyte-specific marker glial fibrillary acidic protein (GFAP, Fig. 2J-L) showed an increased number of astrocytes or astrocytosis and fibrillary processes per astrocytes or astrogliosis surrounding the region of cryptococcal-induced encephalomalacia and GXM accumulation (Fig. 2K-L). Astrocytes also appear necrotized and unable to form a barrier between the damaged brain tissue and the remaining tissue (Fig. 2L). These data indicated that mice infected in bregma develop cerebral cryptococcosis characterized by encephalomalacia near the septum pellucidum, GXM accumulation, and glial activation in the marginal zone of the affected tissue.

The cerebral cortex is one of the brain regions affected by CME (32, 38). Thus, we characterized the progression of cerebral cryptococcosis in mice infected with the fungus in the cortex (Fig. 3). As shown, a coronal section from a cortically infected mouse brain removed at 7-dpi displayed a large cryptococcoma in the ipsilateral cerebral cortex (Fig. 3A; black square), smaller disseminated brain-lesions filled with cryptococci, and signs of ischemia including widespread microinfarcts in deep tissue including the septal nucleus, striatum, and piriform cortex. High magnification images show a cryptococcoma characterized by liquefactive necrosis (Fig. 3B), presence of vacuolated or hueco cells, and the remaining tissue in the margins displaying severe degenerative changes with no inflammatory infiltrate (Fig. 3C). The contralateral cerebral cortex (green square in Fig. 3A) evinced severe interstitial edema forming status spongiosis in the brain parenchyma (Fig. 3D). The leptomeninges (red square in Fig. 3A) showed edema and phagocytic cell recruitment particularly in the pia matter and arachnoid meninges (Fig. 3E). Subarachnoid hemorrhage was observed (blue square in Fig. 3A) and extravasated blood accumulated (white arrows) between pia matter and arachnoid meninges (Fig. 3F). IHC demonstrated ubiquitous ipsilateral and contralateral distribution of GXM with more prominence in the margins of infarction (Fig. 3G-I). Iba-1 staining displayed activated microglia with many and thick ramifications in response to tissue damage due to cryptococcal colonization (Fig. 3J-L). Dystrophic or necrotic microglia can be observed in tissue surrounding the region of expanded encephalomalacia (Fig. 3L). Substantial astrocytosis and astrogliosis was shown surrounding the cryptococcal brain lesion and clumping in cortical tissue (Fig. 3M-O). Our findings indicate that mice i.c. infected in the cortex developed local and scattered cryptococcomas characterized by encephalomalacia, edema, ischemic signs including blood vessel infarcts, meningeal bleeding, significant CPS accumulation, and glial morphological changes.

**Fig. 3.**
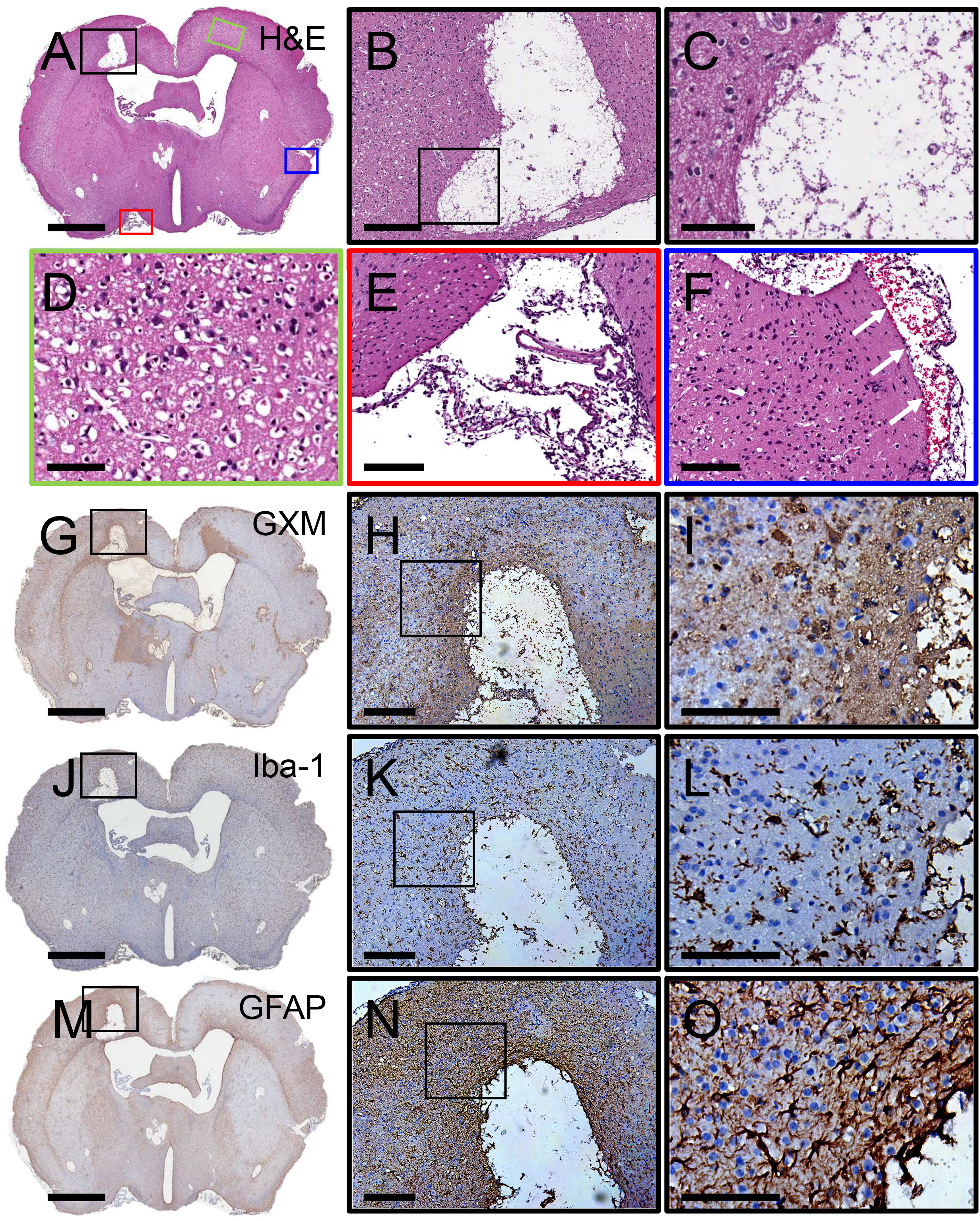
Cerebral cryptococcosis in mice infected in cortical tissue. Representative coronal brain sections (scale bar: 1 mm) of cortical-infected mice stained with **(A-C)** H&E, **(G-I)** GXM-specific mAb 18B7, **(J-L)** Iba-1 (microglia), and **(M-O)** GFAP (astrocytes) are shown after tissue excision 7-dpi. **(D)** Green, **(E)** red, and **(F)** blue boxes are a magnification (scale bar: 100 μm) of specific regions of the brain in **A**. White arrows in **F** show hemorrhage in the subarachnoid space. Middle (**B**, **H**, **K**, **N**; scale bar: 100 μm) and right (**C**, **I**, **L**, **O**; scale bar: 20 μm) panel images are a magnification of the smaller box in the corresponding left-stained section to better show the tissue morphology, GXM distribution (brown stain), and glial cell recruitment/activation (brown stain). Five mice per infection route were evaluated to confirm all the histopathological observations reported in these studies.

In post-mortem brain tissue from CME patients, cryptococci are usually distributed into the corpus striatum of the basal ganglia, where dopaminergic neurons are abundant (32). Given the importance of this region in human disease, we documented the advancement of the cryptococcal infection in the striatum of C57BL/6 mice 20-dpi. A representative brain coronal section at midpoint of the infundibulum displayed aggressive cryptococcoma expansion from the striatum to include the thalamus, hypothalamus, habenula, the microinfarction in contralateral thalamus and hippocampus and frontoparietal cortex (Fig. 4A). The area of ischemic or liquefactive necrosis displayed hemorrhage (Fig. 4B-C), vacuolated macrophages or hueco cells (black arrowheads) and foreign body giant cells (red arrow; Fig. 4D). We observed leptomeningitis with infiltration of activated epithelioid or macrophage-like cells (red square in Fig. 4A; black arrow; Fig. 4E). Perivascular edema and ischemic or angular neurons (white arrows) developed in the contralateral frontoparietal and visual cortex suggesting obstruction of venous drainage system (blue square in Fig. 4A; Fig. 4F). GXM staining the margins of the cryptococcoma, extending to the frontoparietal cortex of the ipsilateral and contralateral sides and the thalamus of the contralateral side (Fig. 4G-I). High magnification images show GXM intensely distributed around the margins of the cryptococcoma and concentrated in the wall of cerebral blood capillaries surrounding the area of encephalomalacia (Fig. 4H-I). Microglia are observed surrounding the cryptococcoma (Fig. 4J). These brain surveillance resident cells are shown activated inside the cryptococcoma (Fig. 4K) and dystrophic surrounding the liquefactive tissue (Fig. 4L). Significant and extensive astrocytosis is shown in the wall of the cryptococcoma that extends to the hippocampus (Fig. 4M). The presence of astrogliosis including astrocytic clumping surrounding the margins of the cryptococcoma is evident (Fig. 4N-O). These results show that direct infection of the basal ganglia is characterized by significant expansion of the cyptococcoma that consists of yeast cells, necrotic tissue with hemorrhage, hueco cells, and foreign body giant cells. Furthermore, the development of meningitis and edema with ischemic neurons can be appreciated especially in brain regions distant from the initial inoculum.

**Fig. 4.**
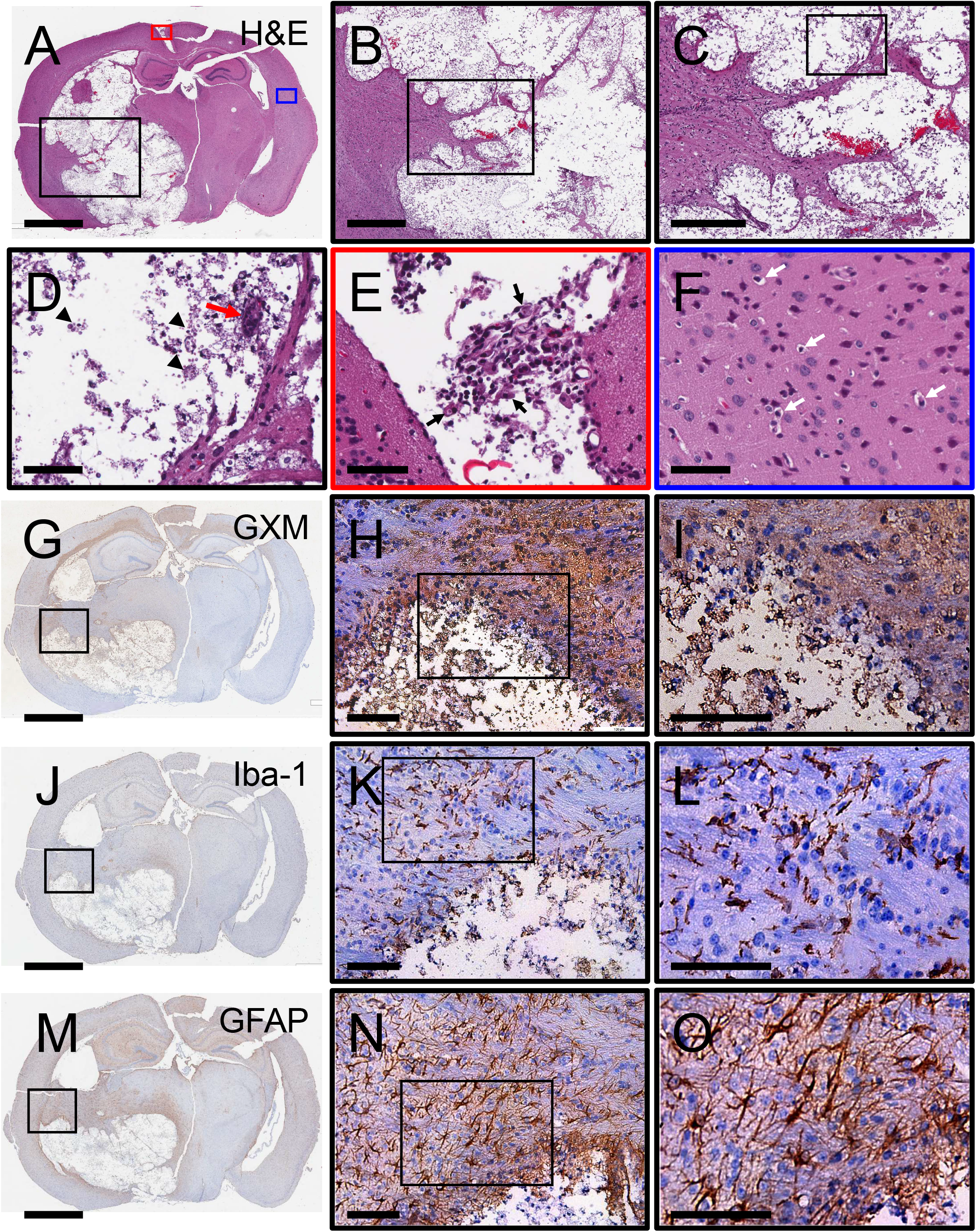
Cerebral cryptococcosis in mice infected in the corpus striatum of the basal ganglia. Representative coronal brain sections (scale bar: 1 mm) of striatal-infected mice stained with **(AD)** H&E, **(G-I)** GXM-specific mAb 18B7, **(J-L)** Iba-1 (microglia), and **(M-O)** GFAP (astrocytes) are shown after tissue excision 20-dpi. **(D)** Arrow heads denote vacuolated macrophages or hueco cells. Red arrow indicates foreign body giant cell. **(E)** Red and **(F)** blue boxes are a magnification (scale bar: 100 μm) of specific regions of the brain in **A**. Black arrow in **E** show activated macrophages or epithelioid cells. White arrows in **F** denote perivascular edema and ischemic neurons. Middle (**B**, **H**, **K**, **N**; scale bar: 100 μm) and right (**C**, **I**, **L**, **O**; scale bar: 20 μm) panel images are a magnification of the smaller box in the corresponding left-stained section to better show the tissue morphology, GXM distribution (brown stain), and glial cell recruitment/activation (brown stain). Five mice per infection route were evaluated to confirm all the histopathological observations reported in these studies.

*C. neoformans* infects the choroid plexus and the ventricles, causing ventriculitis and/or hydrocephalus. This condition is associated with increased ICP (39, 40). Lack of CSF reabsorption in the arachnoid granulations due to the accumulation of cryptococcal polysaccharide and/or yeast cells has been related to increased ICP (41, 42). The presence of CPS in the CSF is used for diagnosing CME in patients, highlighting the importance of understanding how the ventricle environment is affected during *C. neoformans* infection. Therefore, we also studied the progression of *C. neoformans* infection in the brain right ventricle of C57BL/6 mice after tissue removal 7-dpi. A coronal section of brain tissue at midpoint of the infundibulum shows ventriculomegaly, infarction in the hippocampus, and substantial subarachnoid hemorrhage (black arrow; Fig. 5A). Minimal cryptococcoma formation was observed in mice infected in the right ventricle. The area of infarction in the hippocampus exhibited liquefactive necrosis (Fig. 5B). A high magnification image demonstrates minimal inflammation and the presence of a megakaryocyte (blue arrow) that is indicative of thrombosis (Fig. 5C). Ventricle-infected tissue demonstrated extensive distribution of GXM in regions such as the cerebral cortex, meninges, basal ganglia, and corpus callosum (Fig. 5D). GXM was concentrated in the frontoparietal cortex and the wall of large blood vessels and capillaries (Fig. 5E-F). Microglia was mostly concentrated in the hippocampus around the infarction area (Fig. 5G-I). In contrast, astrocytes were distributed all over the brain tissue with astrogliosis mostly observed near blood vessels (Fig. 5J-L). Our observations in ventricle-infected mice demonstrate minimal cryptococcoma formation, minimal inflammation, localized infiltration of microglia, and significant accumulation of GXM especially in blood vessels, which are surrounded by reactive astrocytes.

**Fig. 5.**
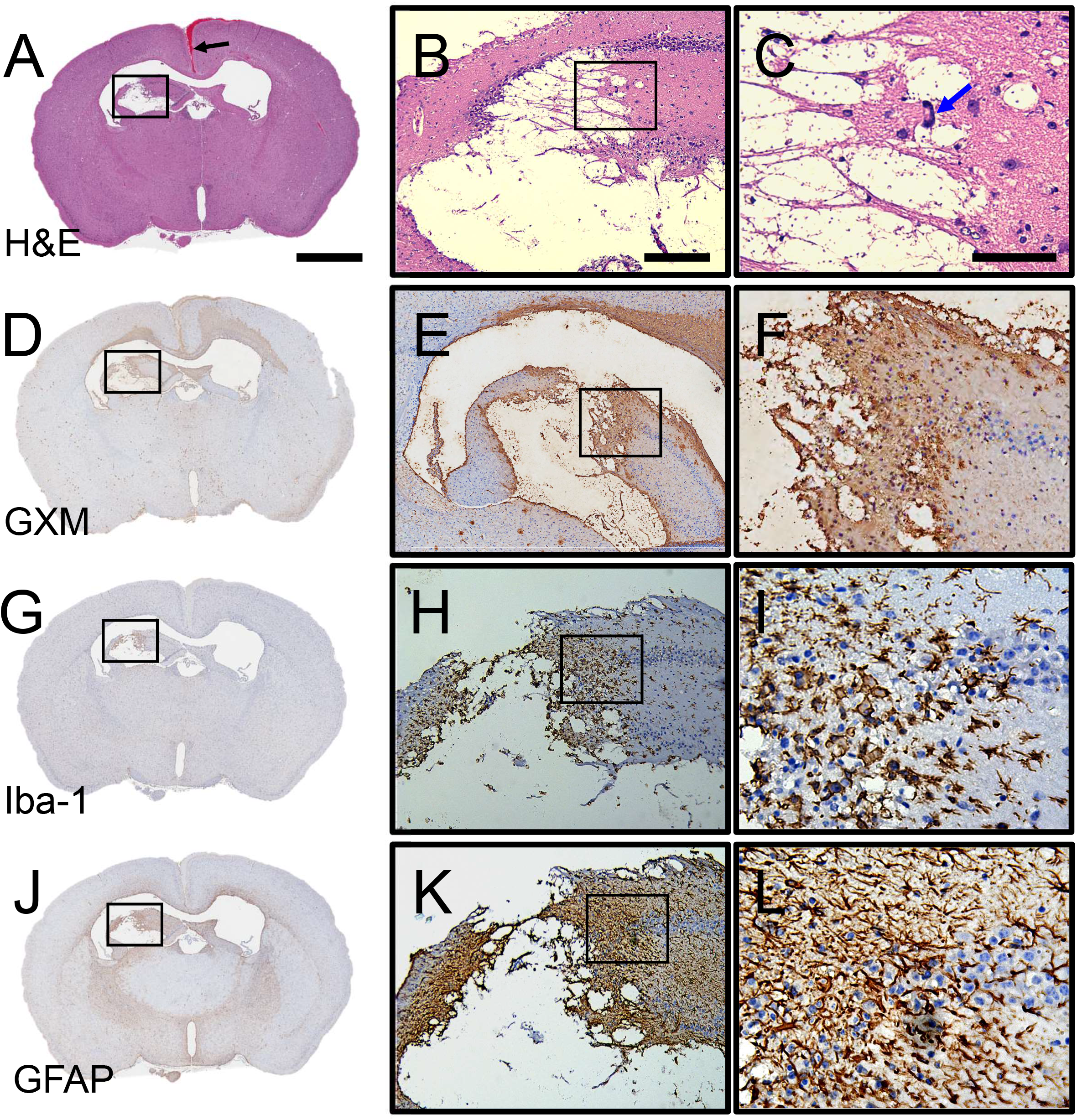
Cerebral cryptococcosis in mice infected in the ventricles. Representative coronal brain sections (scale bar: 1 mm) of ventricle-infected mice stained with **(A-C)** H&E, **(D-F)** GXM-specific mAb 18B7, **(G-I)** Iba-1 (microglia), and **(J-L)** GFAP (astrocytes) are shown. Mouse brains were excised 7-dpi. Middle (scale bar: 100 μm) and right (scale bar: 20 μm) panel images are a magnification of the smaller box in the corresponding left-stained section to better show the tissue morphology, GXM distribution (brown stain), and glial cell recruitment/activation (brown stain). Black arrow in **A** denotes subarachnoid space hemorrhage. Blue arrow in **C** indicates the presence of megakaryocytes and indicative of thrombosis. Five mice per infection route were evaluated to confirm all the histopathological observations reported in these studies.

### Relationship between *C. neoformans* GXM localization and glial cell distribution in the mouse brain

We analyzed the intensity correlation of either Iba-1 and GFAP markers relative to *C. neoformans* GXM in coronal tissue sections excised from mice infected in the bregma, cortex, striatum, or ventricles using the NIH Image J software (Fig. 6). Mice infected in bregma showed positive correlations between Iba-1 (R^2^ = 0.3533; *P* = 0.0152; Fig. 6A) and GFAP (R^2^ = 0.2942; *P* = 0.0299; Fig. 6B) stained cells and GXM distribution. Iba-1 and GXM (R^2^ = 0.1269; *P* = 0.1468; Fig. 6C) staining did not correlate in mice infected in the cerebral cortex, although there was a correlation between the GFAP and GXM (R^2^ = 0.2472; *P* = 0.0358; Fig. 6D) intensity. In addition, mice infected in the striatum evinced positive correlations between the intensity of Iba-1 (R^2^ = 0.5915; *P* = 0.0035; Fig. 6E) and GFAP (R^2^ = 0.4958; *P* = 0.0106; Fig. 6F) staining and the intensity of GXM in tissue. There was no correlation between Iba-1 (R^2^ = 0.0009; *P* = 0.8915; Fig. 6G) and GFAP (R^2^ = 0.0671; *P* = 0.2215; Fig. 6H) intensity and GXM in ventricle-infected mice. These analyses demonstrate that glial cell responses to GXM accumulation are different depending on the localization of the *C. neoformans* infection in the brain.

**Fig. 6.**
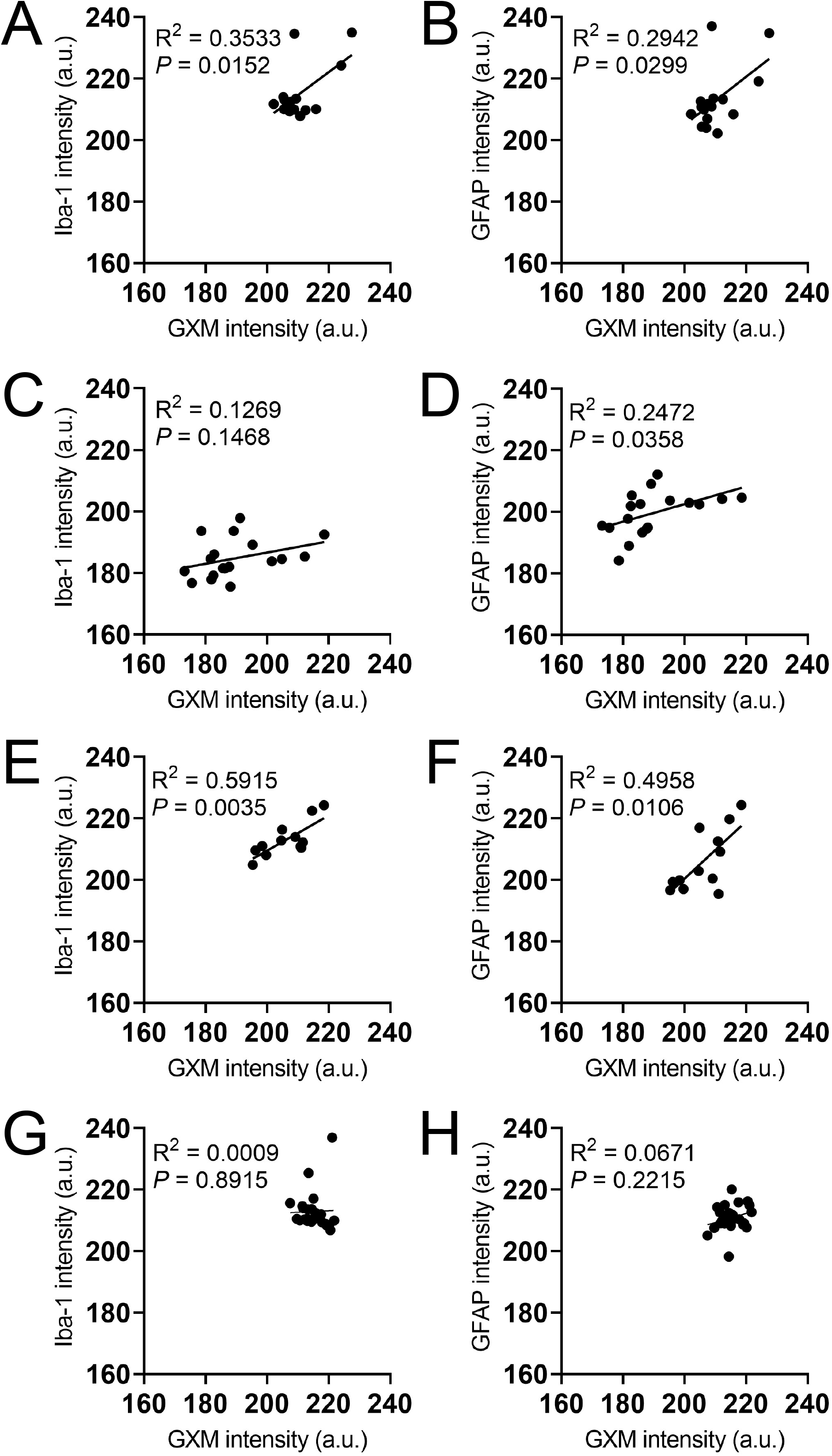
Glia distribution and GXM localization correlation vary depending on the brain region infection. Correlations between GXM localization and glial cell infiltration in brain tissue of mice infected with *C. neoformans* strain H99 cells in **(A-B)** bregma, **(C-D)** cortex, **(E-F)** striatum, and **(G-H)** ventricles. GXM, microglia, and astrocytes were stained with mAb 18B7, Iba-1, and GFAP, respectively. The intensity of each marker was analyzed in coronal tissue sections using the NIH Image J software. The R^2^ and *P* values for each regression are also indicated.

### Mortality is associated with GXM localization and hemorrhage in the subarachnoid space

Approximately 30% of clinical cases have reported infarcts and blood vessel damage in patients with severe CME (43). Histopathological examinations of mice infected i.c. with *C. neoformans* at the time of death demonstrated that mortality was associated to hemorrhage in the brain subarachnoid space and GXM accumulation in this tissue area (Fig. 7). Brain coronal tissue section at midpoint of the infundibulum from an uninfected and healthy mouse showed normal anatomy and normal leptomeninges covering the retrosplenial and frontoparietal cortex (Fig. 7A-B). Mice infected in bregma (Fig. 7C), and brain tissue removed 7-dpi displayed notable subarachnoid hemorrhage (black arrows; Fig. 7D) and moderate GXM deposition in the glia limitans and meninges (red arrows; Fig. 7E). Cortical tissue excised 7-dpi demonstrated marked hemorrhage in the subarachnoid space (Fig. 7F-G), bulging of the superior sagittal sinus (black arrow; Fig. 7F-G), and extensive GXM distribution in the leptomeninges especially near the bleeding (red arrows; Fig. 7H). Moreover, mice infected with the fungus in the striatum (Fig. 7I-K) or ventricles (Fig. 7L-N) also showed considerable subarachnoid space and superior sagittal sinus bleeding (black arrows; Fig. 7J and M). GXM was profusely dispersed in the leptomeninges (red arrows; Fig. 7K and N). Our examinations suggest that mice mortality may be related to extensive *C. neoformans* CPS localization near the subarachnoid space blood vessel infarcts and hemorrhage.

**Fig. 7.**
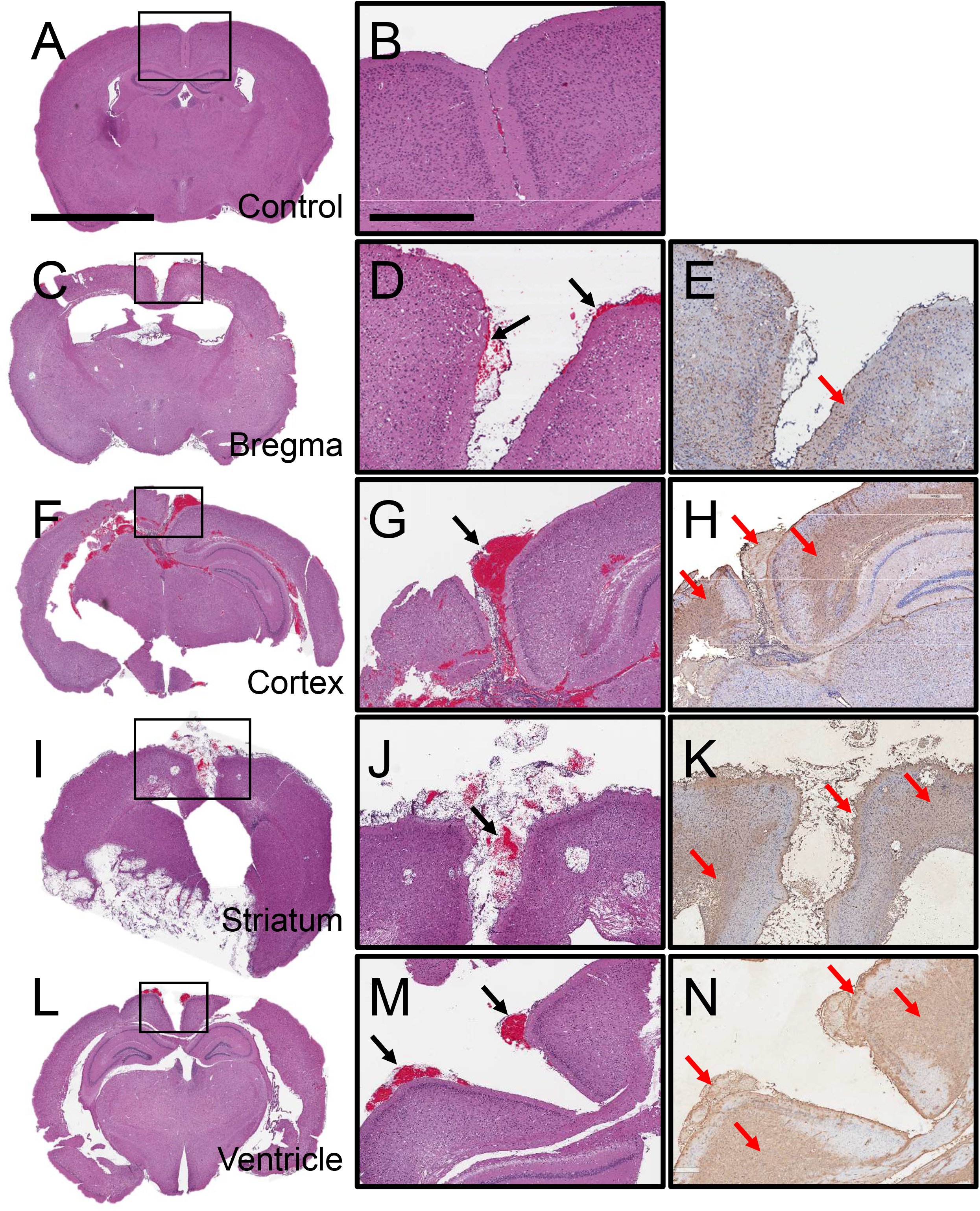
*C. neoformans* GXM localization is associated with subarachnoid space hemorrhage in i.c. infected mice. Representative coronal brain sections (scale bar: 2 mm) of **(A-B)** uninfected (control) and **(C-E)** bregma-, **(F-H)** cortex-, **(I-K)** striatum-, and **(L-N)** ventricle-infected mice with *C. neoformans* H99 strain. Each brain section was stained with H&E and GXM-binding mAb 18B7. Uninfected (control) and infected mice in bregma, cortex, and ventricle were euthanized 7-dpi. Striatum-infected rodents were sacrificed 20-dpi. Middle (H&E) and right (GXM-labeled; brown stain) panel images (scale bar: 200 μm) are a magnification of the smaller box in the corresponding left H&E-stained section to better show the tissue morphology and GXM distribution. Black arrows indicate subarachnoid space hemorrhage. Red arrows demonstrate GXM accumulation in the subarachnoid space. Five mice per infection route were evaluated to confirm all the histopathological observations reported in these studies.

### *C. neoformans* cortical infection results in apoptosis of neurons in the hippocampus

We further investigated the sequalae of cerebral cortical *C. neoformans* infection in C57BL/6 mice. Histopathological examination of the hippocampus, one of the main brain regions associated with memory and cognitive function, after fungal infection of the cerebral cortex showed severe edema and passive hyperemia likely due to obstruction of cerebral blood flow (Fig. 8A). We also observed signs of neurodegeneration in the dentate gyrus, appearing as cytoplasmic vacuolation. High power magnification (white rectangle box) also indicated apoptotic pyramidal neurons (red arrows) distributed across the CA3 region (Fig. 8B), suggesting the possibility that cerebral cryptococcosis, particularly cortical colonization, may cause altered behavior in mice or altered mental status in patients with CME.

**Fig. 8.**
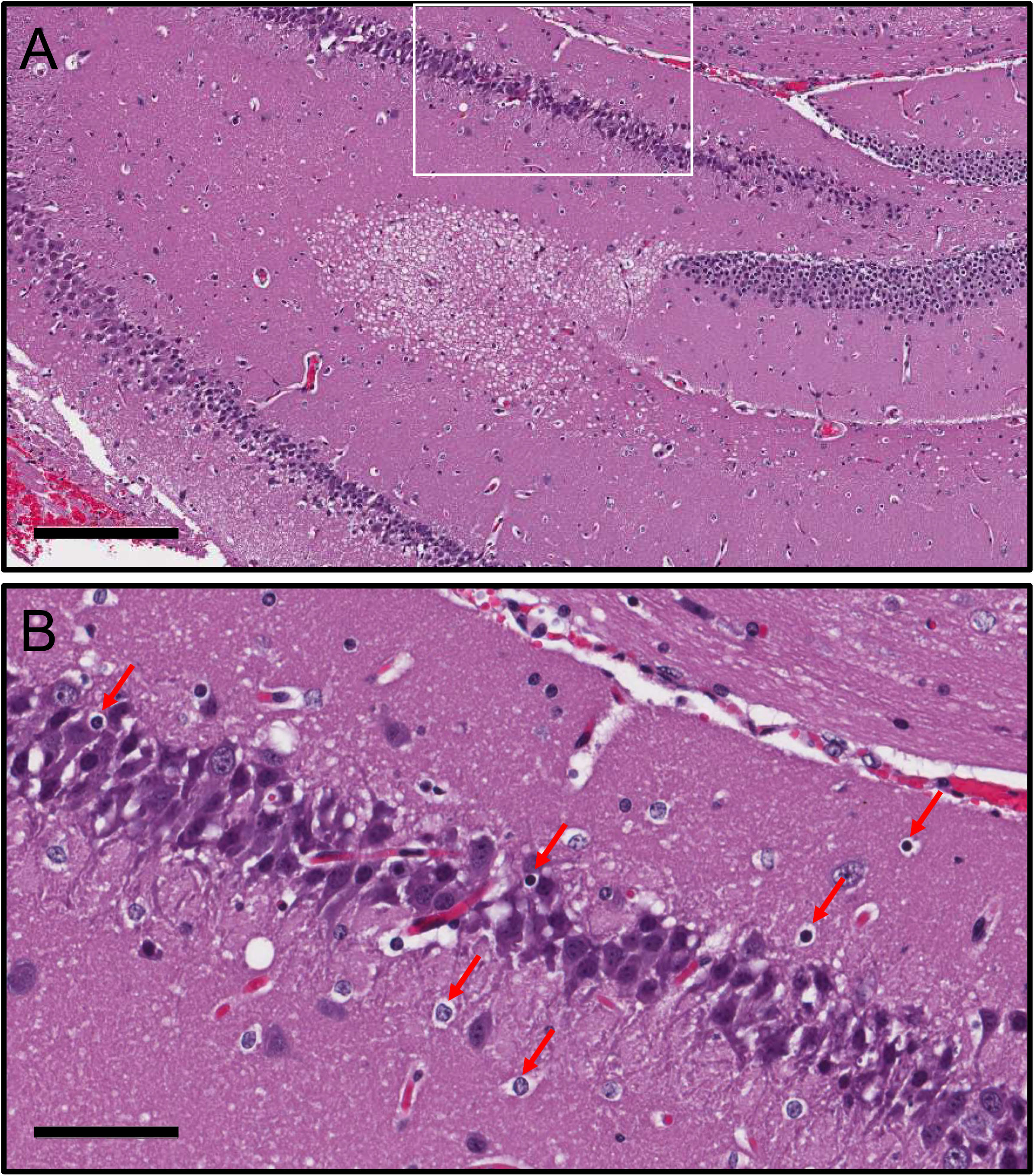
*C. neoformans* cortical infection progresses to hydrocephalus and causes neuronal apoptosis in the hippocampus. **(A)** Representative hippocampal section stained with H&E. Scale bar: 200 μm. **(B)** Magnification of the white rectangle in **A.** Red arrows show apoptotic neurons in the CA3 region of the hippocampus. Scale bar: 60 μm. Five mice per infection route were evaluated to confirm all the histopathological observations reported in these studies.

### *C. neoformans* i.c. infection causes microinfarctions and tissue bleeding

We documented the effects of *C. neoformans* i.c. infection in brain tissue (Fig. 9). Hemorrhagic microinfarctions were observed in the frontal cortex (rectangle; Fig. 9A-B) and increased infiltration of gitter cells (blue arrows; Fig. 9C), which are enlarged phagocytic cells of microglial origin having the cytoplasm distended with lipid granules. Prussian blue staining confirmed hemosiderosis or excessive accumulation of iron deposits that arises from hemorrhage (Fig. 9C; inset). Hemorrhagic microinfarctions were visualized near to cryptococcomas in the corpus callosum (Fig. 9D). We observed in this region, leukomalacia or necrosis of the white matter, degeneration and demyelination of the nerve fibers, and recruitment of gitter cells (Fig. 9E) in the presence of erythrocytes (black arrows; Fig. 9F). Hemorrhage in the margins of a cryptococcoma (rectangle box; Fig. 9G) shows extensive aggregation of gitter cells around the bleeding area and cryptococcal brain lesion (Fig. 9H). High power magnification confirmed significant gitter cell recruitment (blue arrows) into the hemorrhage region (Fig. 9I). i.c. *C. neoformans* infection of the striatum results in extensive encephalomalacia, multiple hemorrhage foci (Fig. 9J), and considerable aggregation of polymorphonuclear cells around the blood vessels and in the lesion (Fig. 9K-L). A coronal section of mice infected with *C. neoformans* show multiple cryptococcomas scattered in the posterior limb of internal capsule, the putamen (black rectangle box), nucleus accumbens, and septal nucleus (Fig. 9M). Microthrombus in the wall of a blood capillary and well circumscribed infarcted tissue from ischemia and liquefactive necrosis (red arrows; Fig. 9N). These images show that cerebral cryptococcosis after i.c. infection causes thrombosis of blood capillaries in the subarachnoid spaces resulting in ischemia, microinfarctions, and hemorrhage in regions of the CNS where there is fungal colonization and adjacent GXM accumulation.

**Fig. 9.**
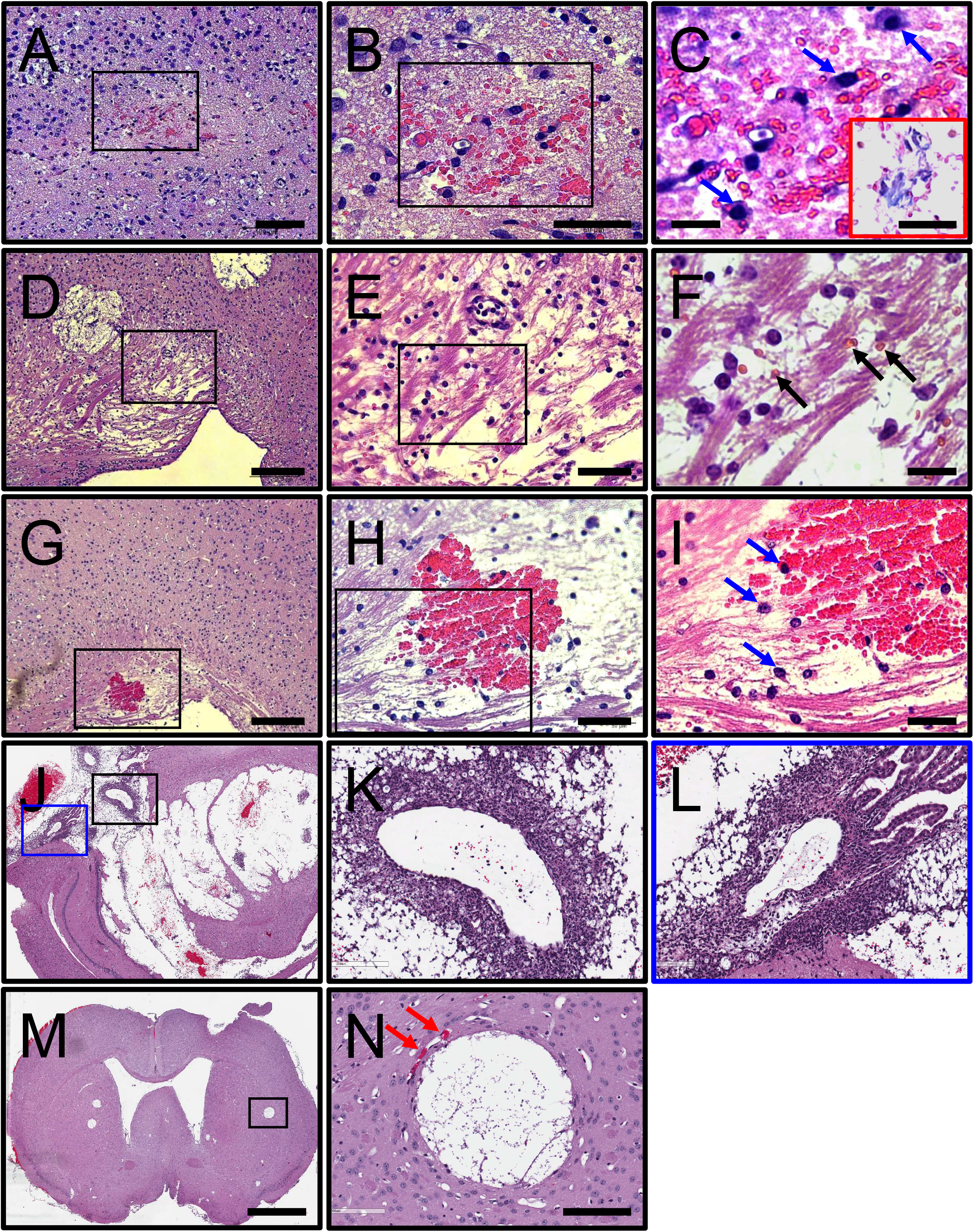
*C. neoformans* infection causes microinfarctions and hemorrhage in cerebral tissue. **(A-B)** Hemorrhagic microinfarction (black rectangles) in the frontal cortex and **(C)** infiltration of gitter cells (blue arrows) from microglia origin 7-dpi. Black rectangles in **A** and **B** correspond to a magnified region of the tissue shown in the right-side panel. Scale bars: **(A)** 100 μm, **(B)** 60 μm, and **(C)** 20 μm. Red inset exhibit Prussian blue staining to confirm that hemosiderosis or focal deposition of iron due to local bleeding Scale bar: 10 μm. **(D)** Leukomalacia or necrosis of white matter near a cryptococcal lesion in the corpus callosum. **(E-F)** Magnified region in **D** exhibits demyelination of the nerve fibers, recruitment of gitter cells, and the presence of erythrocytes (black arrows). Scale bars: **(D)** 200 μm, **(E)** 50 μm, and **(F)** 20 μm. **(G-I)** Extensive bleeding, degeneration of nerve fibers, and gitter cell infiltration (blue arrows) in the corpus callosum. Scale bars: **(D)** 200 μm, **(E)** 50 μm, and **(F)** 20 μm. **(J)** i.c. *C. neoformans* infection of the striatum results in extensive encephalomalacia, multiple hemorrhage foci, and **(K-L)** considerable aggregation of polymorphonuclear cells around the blood vessels and in the lesion. **(K)** Black and **(L)** blue boxes in **J** are magnification of those regions. **(M)** Coronal section at the level of anterior commissure in the right ventricle of mice infected with *C. neoformans*. Scale bar: 1 mm. **(N)** Magnification of a cryptococcoma (black rectangle box) showing microthrombus in the wall of blood capillary, resulting in microinfarctions (red arrows). Scale bar: 200 μm.

### *C. neoformans* GXM extensively deposits in the vasculature

*C. neoformans* GXM is extensively released and accumulates in the CSF and brain tissue during infection. Recently, it was shown in zebra fish that blood vessel occlusion by this encapsulated fungus is a mechanism for hemorrhagic dissemination of infection (44). Given that GXM is associated with excessive bleeding in the subarachnoid space and increased mortality in mice i.c. infected, we examined the localization of CPS in brain tissue with emphasis in the vasculature. GXM IHC with mAb 18B7 of a coronal brain section of a mouse infected in the right ventricle with *C. neoformans* showed dispersed polysaccharide throughout the tissue and in the wall of blood vessels in the cerebral cortex, hippocampus, habenula, thalamus and hypothalamus (Fig. 10A). Considerable GXM deposition was visualized in the wall of blood capillaries (red arrow) and large vessels (black rectangle box; Fig. 10B) in the thalamus. Higher magnification image demonstrated significant GXM deposition (brown stain) in the walls of a medium size blood vessel and several surrounding capillaries (Fig. 10C). Furthermore, GXM accumulated mainly in the wall of large blood vessels in the subarachnoid space (red arrows; Fig. 10D). Our findings that the fungal CPS localizes in the vasculature provides supportive evidence that GXM and cryptococci may accumulate and obstruct blood vessels in the CNS resulting in microinfarctions, hemorrhage, and exacerbating death.

**Fig. 10.**
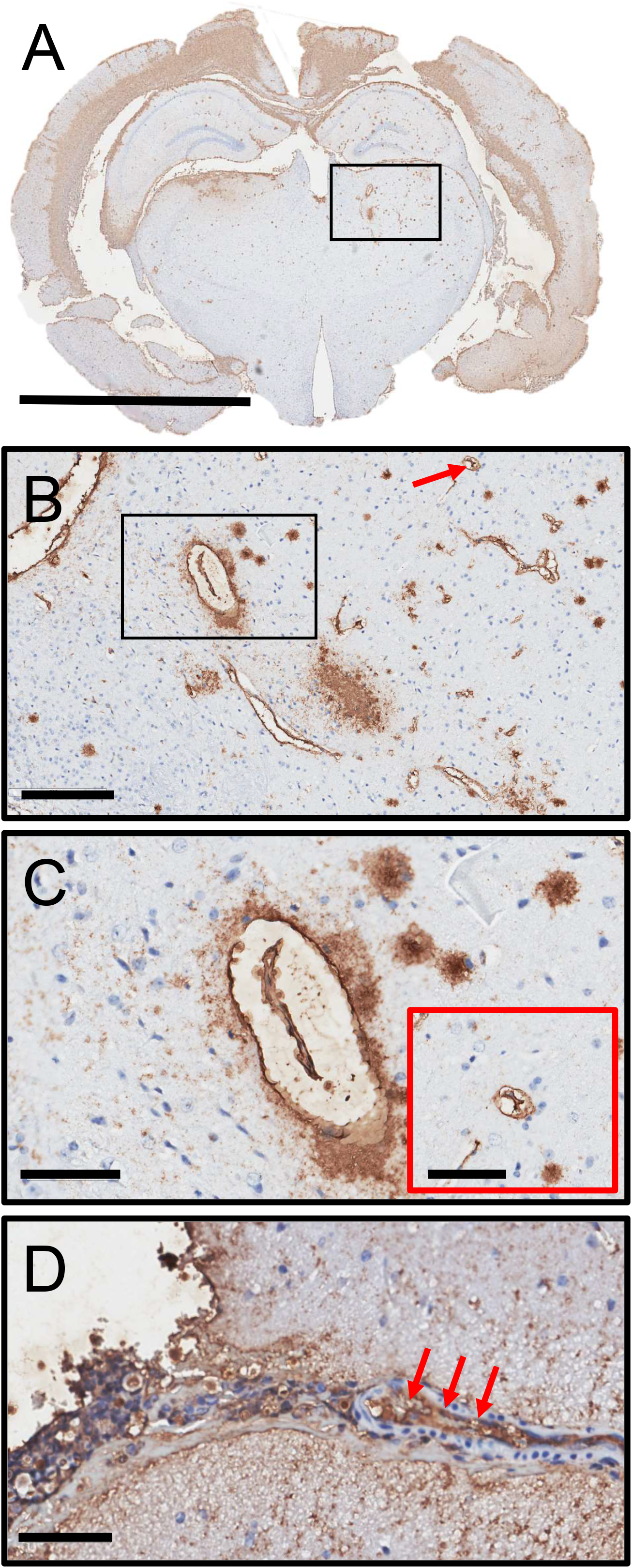
*C. neoformans* GXM abundantly localizes in the vasculature. **(A)** Representative GXM immunohistochemistry (IHC) with mAb 18B7 of a coronal brain section of a mouse infected in the right ventricle with *C. neoformans*. Scale bar: 2 mm. **(B)** Magnification of the small box in **A** shows specific localization of GXM in capillaries (red arrow) and blood vessels (black rectangle) of the thalamus. Scale bar: 200 μm. **(C)** Magnification of the black rectangle in **B** demonstrates substantial GXM deposition (brown stain) in the wall and the surroundings of a medium size blood vessel and several capillaries. Scale bar: 60 μm. Inset shows capsular polysaccharide around a capillary. Scale bar: 10 μm. **(D)** Extensive accumulation of GXM in the wall of an artery (red arrows) near the subarachnoid space. Scale bar: 60 μm.

### Mice infected with *C. neoformans* i.c. develop ischemic necrosis

Histopathological examinations of the frontal cortex from C57BL/6 mice infected with *C. neoformans* demonstrate noticeable ischemic necrosis of the gray matter (black box rectangle; Fig. 11A). Ischemic necrosis is consistent with the presence of clear spaces of liquefactive tissue where the damaged neurons and tissue are phagocytosed and then removed. High power magnification shows infiltration and accumulation of gitter cells (blue arrows) with foamy cytoplasm that results from engulfing necrotic tissue (Fig. 11B). In addition, microinfarctions were observed in cortical tissue (red arrows; Fig. 11C). A *C. neoformans* lesion or cryptococcoma in the striatum of mice displayed a foreign body giant cells (black rectangle) surrounded by fungal cells and numerous phagocytic cells with vacuolated cytoplasm (Fig. 11D-F). Damage of cerebral capillaries (encircled red areas) and microthrombus in the striatum of infected mice are shown with presence of a few Langhans giant cells (red arrows) and little macrophagic infiltration indicating poor granuloma formation (Fig. 11G). Also, coagulative necrosis is observed in neurons with pyknotic nuclei and increased cytoplasmic acidophilia. There were no epithelioid cellular responses, which are characteristic for granulomatous inflammation. There is noticeable accumulation of cryptococci in liquefactive tissue around the border of the brain lesion likely waiting for more liquefaction of tissue to proliferate further. Our results show that *C. neoformans* cerebral colonization causes extensive vascular damage and ischemic necrosis, two observations associated with mice mortality.

**Fig. 11.**
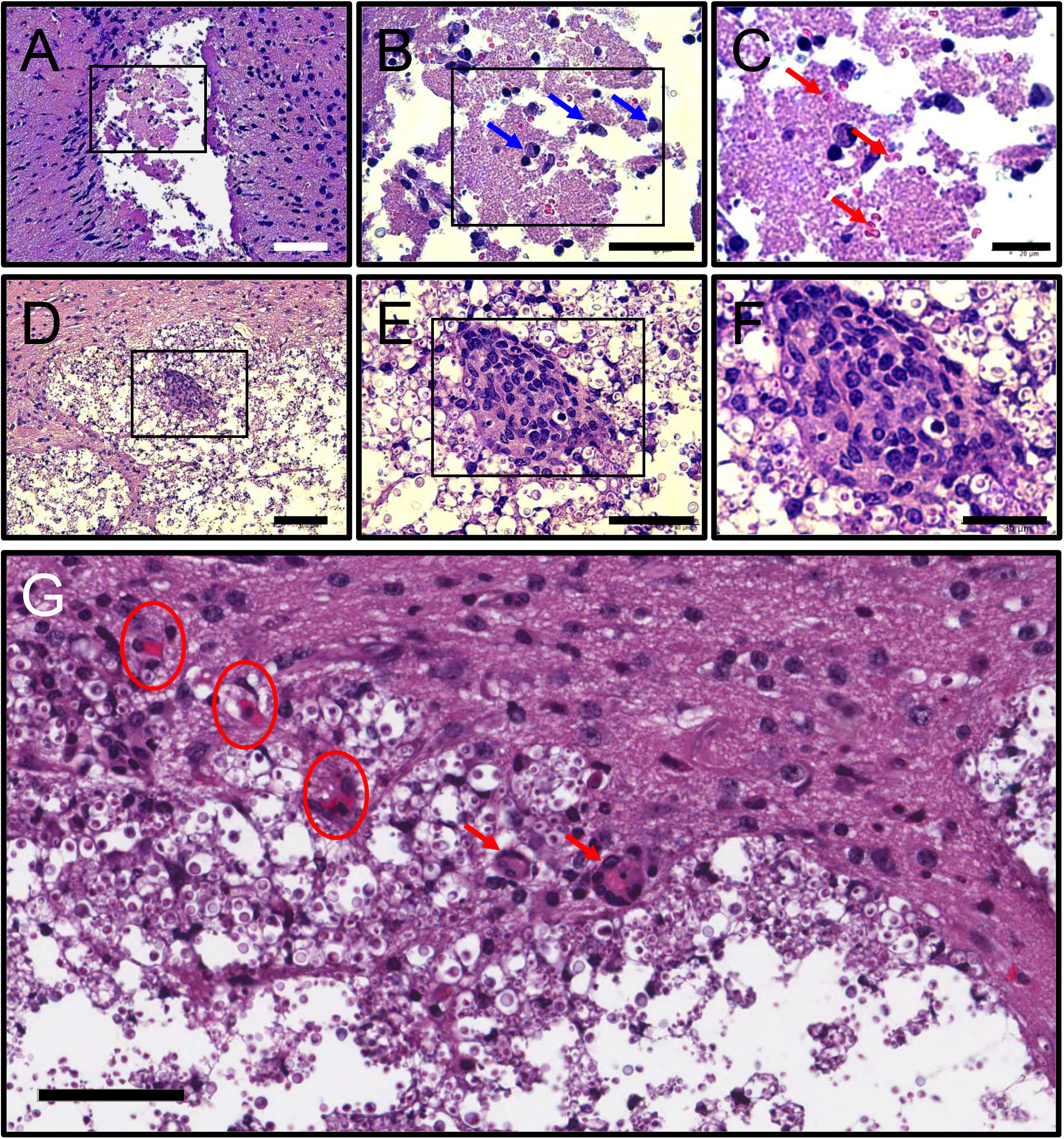
C57BL/6 mice infected with *C. neoformans* i.c. develop ischemic necrosis. H&E staining of the frontal cortex showing **(A)** necrosis of the gray matter, **(B)** accumulation of gitter cells (blue arrows in magnified rectangle) with foamy cytoplasms from engulfing necrotic tissue, and **(C)** microinfarctions (red arrows). Scale bars: **(A)** 100 μm, **(B)** 50 μm, and **(C)** 20 μm. **(D-F)** *C. neoformans* lesion or cryptococcoma in the striatum of mice show foreign body giant cells (black rectangle) surrounded by fungal cells and numerous phagocytic cells with vacuolated cytoplasm. Scale bars: **(A)** 100 μm, **(B)** 90 μm, and **(C)** 30 μm. **(G)** Damage of cerebral capillaries (encircled areas) and microthrombus in the striatum of infected mice. Langhans giant cells are depicted by red arrows indicating poorly formed granulomas in striatal tissue. Scale bar: 60 μm. Five mice per infection route were evaluated to confirm all the histopathological observations reported in these studies.

### Microglia inside of a cryptococcoma becomes activated while combatting the infection

We documented the microglial responses to *C. neoformans* i.c. infection in the striatum (Fig. 12). Histopathology showed phagocytic cells engulfing cryptococcal cells and necrotic tissue in the cryptococcoma (Fig. 12A). The presence of free yeast cells in the brain lesion and phagocyte recruitment to the damage tissue or the area of encephalomalacia is evident (Fig. 12A). IHC for Iba-1 in the cryptococcoma displays microglia or hueco cells with vacuolated cytoplasm (red arrows) responding to the progressive encephalomalacia (Fig. 12B). We identified engulfed cryptococci by several microglia in the cryptococcoma lesion. These images show that microglia or hueco cells are one of the hallmarks of a cryptococcoma. They activate and try to fight the infection by this encapsulated fungus while removing necrotic or damaged tissue.

**Fig. 12.**
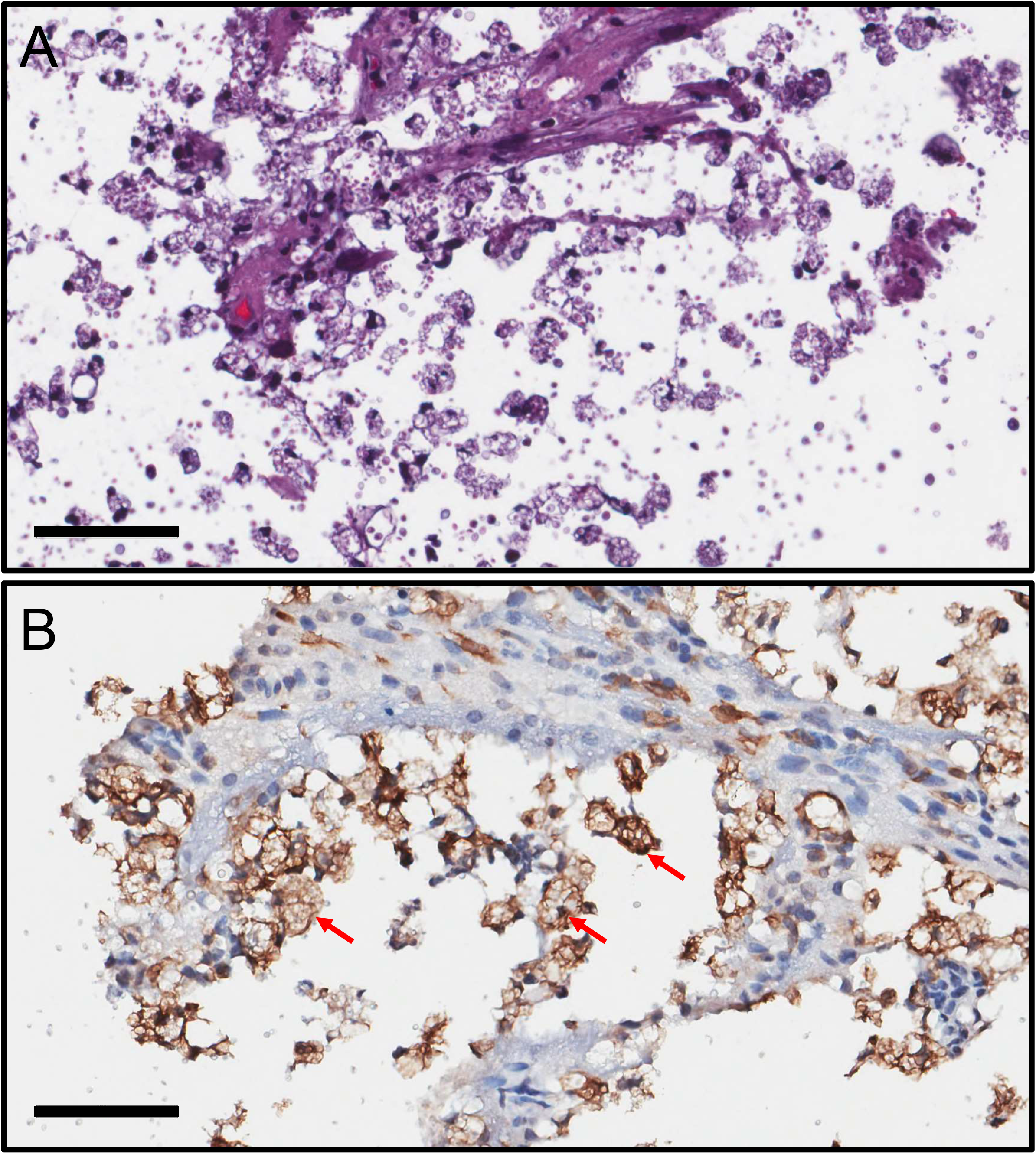
Activated microglia inside of a cryptococcoma fighting the infection in the striatum. **(A)** H&E straining demonstrating phagocytic cells engulfing fungal cells and necrotic tissue in the cryptococcoma. **(B)** IHC for Iba-1 in the cryptococcal lesion shows microglia or hueco cells with vacuolated cytoplasm (red arrows) responding to the progressive encephalomalacia. For **A** and **B**, scale bar: 60 μm.

## DISCUSSION

CME is the most important manifestation of *C. neoformans* infection. Most of our understanding of neurocryptococcosis comes from post-mortem observations in humans (32, 38). One of the main roadblocks to study CME is the availability of well-characterized or reliable animal models where the disease progression and dynamics can be systematically studied, and the pathological manifestations show similarities to those presented in humans. Although not a natural route of *C. neoformans* infection, we used the i.c. model to study the development of CME and the cellular/physiological changes/responses in specific regions of the brain with a controlled inoculum (10^4^ fungi) and a suitable volume (1 μL). Stereotaxic injections, although invasive, are extensively used in neuroscience research to gain knowledge of CNS infectious disease processes. In fact, tissue damage/response from the injection can be easily controlled by using saline-injected and naïve animals. In addition, i.c. infection models have been reported in multiple diseases including cryptococcosis (45–48), although not clearly described in studies with *C. neoformans*. An advantage of the i.c. *C. neoformans* model of infection over the intranasal or intratracheal (pulmonary) and intravenous (systemic) models is that the fungal brain dissemination number and the brain region location of infection can be controlled. This is important because the effect and progression of the cryptococcal infection in specific regions of the brain can be characterized. Furthermore, the impact of CME on behavior alteration including motor and cognition can be simply examined. In contrast, the need of mouse-specialized surgical expertise, basic to intermediate knowledge of neuroscience, and the availability of specific equipment such as a stereotaxic apparatus are perhaps the main disadvantages.

Cerebral cryptococcosis in humans is characterized by meningoencephalitis with (cryptococcoma) and without (no cryptococcoma) involvement of the brain parenchyma, especially in immunosuppressed and immunocompetent individuals, respectively (32). There are multiple mechanisms proposed that can lead to meningitis including rupture of cryptococcomas and spread of infection from the perivascular space to the CNS (10, 49–51). Mice infected in the cortex and ventricles with *C. neoformans* showed early mortality, although they exhibited different disease development. Rodents infected in cortical tissue demonstrated cryptococcoma formation, hydrocephalus, edema, and subarachnoid hemorrhage while microinfarctions were minimal. Meningitis in these mice was prominent considering that the meninges directly cover the cortex, and cryptococci and GXM easily reach these membranes through circulation. Similarly, mice infected in the striatum displayed comparable pathology to those infected in the cortex, with the exception that they began dying approximately a week later despite development of microinfarctions. The last mouse died 12 days after those infected in cortical tissue. It is plausible that infection in the striatum, being in a deep region of the brain, takes longer for the fungus to first colonize and form a cryptococcoma, release sufficient GXM that impairs the migration and effector functions of microglia and other inflammatory cells, and cause tissue necrosis. It is likely that free yeast cells or yeast cells engulfed by microglia escape the biofilm-like lesion and enter the capillary network of the CNS, reaching the subarachnoid space and resulting in meningitis and hemorrhage. In this regard, we found similar fungal loads and brain weights in all the routes of infection, validating this possibility. In contrast, intrathecal infection in the ventricles evinced comparable pathologies as those documented in mice infected in the cortex or striatum without brain parenchyma involvement or no cryptococcoma formation. When *C. neoformans* is injected into the ventricles and directly delivered into the CSF, cryptococci reach the meninges and subarachnoid space, rapidly killing the mice (at ~8-dpi). Although there is no cryptococcoma formation like in humans (52), GXM diffuses profusely throughout the tissue, showing affinity for blood vessels where the CPS accumulates (53). It would be interesting to infect the mice intrathecally with a lower inoculum (10^3^ cryptococci) to determine if survival is prolonged to study the infection longer and still have no brain parenchyma involvement. It is apparent from our studies that the region of cryptococcal CNS access as well as the colonization process are important in the progression of CME including the live or death outcomes in patients. Our studies demonstrate, both, cortex and striatum, routes of cryptococcal i.c. infection represent acceptable models to study CME with brain tissue involvement whereas ventricular infection is appropriate to study cerebral cryptococcosis limited to the subarachnoid space.

Individuals with HIV^+^ and CME show little inflammation in the CNS due to their reduced number of CD4^+^ T cells (32). Our findings demonstrated that regardless of the location of the brain infection, cryptococcal infection shows minimal inflammation independent of whether there is tissue or just subarachnoid space involvement. Given that *C. neoformans* extensively releases its GXM that accumulates expansively around cryptococcomas or throughout brain tissue, this CPS inhibits inflammatory responses including recruitment and effector functions such as phagocytosis particularly by microglia (54), the resident cells of immunity in the CNS. GXM inhibits neutrophil migration to microglial-produced IL-8 (55) and the rolling and attachment of these leukocytes to endothelial cells by interfering with their adhesion molecules (56, 57). *C. neoformans*’ neurotropism may be related to the inability of the host in mounting an effective neuroimmune response against the fungus, particularly in immunosuppressed individuals. Lee and colleagues previously suggested that the impaired immunity in patients with AIDS promotes the accumulation of mostly extracellular *C. neoformans* within the brain (32), and that deficient macrophage/microglial effector function may be responsible for the altered pathology. Our histological data show ischemia in tissue with cryptococcomas that was characterized by the recruitment and accumulation of gitter cells with foamy cytoplasms that result from engulfing necrotic tissue and considerable vascular damage. Ischemic stroke is a recognized complication of CME in the acute phase and is thought to be mediated by an infectious vasculitis (58). *C. neoformans* can also induce extensive fibrosis of the subarachnoid space, which may compress small veins mechanically inducing venule congestion and massive cerebral infarction (59). We documented proliferation and accumulation of cryptococci in liquefactive tissue around the border of the brain lesion and considerable deposition of GXM in blood vessels, which may cause compression and occlusion resulting in thrombosis and microinfarctions. In this regard, a recent study on zebra fish demonstrated that cryptococci become trapped in blood vessels, proliferate, cause vascular damage, cortical microinfarctions, and hemorrhage during CME (44). Structural studies on blood vessels and endothelial cells suggest that *C. neoformans* and possibly GXM cause alterations of tight junction and adhesion proteins impairing cell-cell interactions at the blood-brain barrier interface and vasodilation, which affect the vascular tension and facilitates fungal dissemination (44, 60).

The role of glia in CME is poorly understood. Post-mortem tissue from humans have shown the association of glial cells and GXM deposition (53). We established that glial cell responses to the accumulation of CPS is dependent on the CNS region infected by *C. neoformans*. High deposition of GXM in bregma and the striatum correlated with the presence of both microglia and astrocytes. GXM accumulation in cortical tissue correlated with astrocyte presence but not with microglia. The deposition of CPS in mice infected in the ventricles did not correlate with astrocytes nor microglia. It is possible that the prolonged survival documented in mice infected in bregma and the striatum is associated with the astrocyte/microglia responses and their communication with each other or with other cells including those recruited from the periphery. For example, CPS immunoreactivity has been documented in activated microglia interacting with cryptococcal lesions or associated with destructive parenchymal lesions such as necrotizing tissue or infarcts (53). Similarly, GXM has been identified inside of microglia and bound to the cell body of reactive astrocytes (53). However, the mechanisms by which the CPS impact glial cell functions are not well-understood. Thus, further investigations are needed to elucidate the importance of a concerted astrocyte/microglia response against *C. neoformans* in the CNS.

In conclusion, we developed and systematically described a murine model to study the dynamics of cerebral cryptococcosis. Our model recapitulates many of the human pathological manifestations depending on the brain region where the fungus is delivered, including the presence of foreign body giant cells, vacuolated microglia or hueco cells, granulomatous meningitis, neuronal necrosis, and accumulation of the CPS in blood vessels. In addition, the pathogenesis of cerebral cryptococcosis progresses differently depending on the region of infection, making these models suitable to study CME with (e.g., cryptococcoma or cryptococcal brain lesions) or without involvement of the brain parenchyma that result in meningitis, edema, hydrocephalus, and subarachnoid hemorrhage leading to death. Investigations adopting this mouse model of cryptococcal infection may assist us in elucidating critical questions associated with this medically important mycosis, as well as the development of new therapeutics and treatments to combat the CNS disease, which is responsible for significant morbidity and mortality.

## MATERIALS AND METHODS

### C. neoformans

*C. neoformans* strain H99 (serotype A) was isolated and kindly provided by John Perfect at Duke University. Yeasts were grown in Sabouraud dextrose broth (pH 5.6; BD Difco) for 24 h at 30°C in an orbital shaker (Thermo Fisher) set at 150 rpm (to early stationary phase).

### i.c. infection with *C. neoformans*, animal care, and survivability studies

All animal studies were conducted according to the experimental practices and standards approved by the Institutional Animal Care and Use Committee (IACUC) at the University of Florida (Protocol #: 202011067). The IACUC at the University of Florida approved this study. The equipment setup for mouse stereotaxic surgeries and i.c. infections are illustrated in the Supplemental Fig. 1. C57BL/6 female mice (6-8 weeks old; Envigo) were anesthetized using isoflurane (3-5% for induction and 1-2% maintenance; model: VetFlo™ Vaporizer Single Channel Anesthesia System, Kent Scientific), placed in prone position over a heating pad (model: RightTemp® Jr., Kent Scientific), and prepped using aseptic techniques. A local anesthetic, bupivacaine or ropivacaine (0.05%), was administered subcutaneously in the incision. The fur on the skull was carefully shaved off and the animal was placed in a stereotaxic apparatus (model: 940; Kopf Instruments). The mouse’s head was secured using the instrument’s ear bars, incisor bar, and nose clamp as is customary for standard stereotaxic procedures. The skin on the shaved head was prepped with 3 alternating, outward spiral applications of 2-4% chlorhexidine scrub and 70% alcohol prior to incision, and a midline 5-mm incision was made using a scalpel under the guidance of a dissecting microscope. The periosteum was reflected open using small surgical clips. Bregma and lambda skull suture junctions were identified using a stereotaxic brain atlas (e.g., The Allen Mouse Brain Atlas; https://mouse.brain-map.org/static/atlas). Using a small hand-held drill (model: Ideal microdrill; Braintree Scientific), the skull was thinned until the underlying dura mater was visible and a micropipette was brought to the correct stereotaxic position and lowered until it touched the exposed dura. The craniotomy was around 1 mm in diameter. The cut tip of the micropipette has a sharp edge, which will allow easy penetration of the dura of mice. A 1-μL suspension containing 10^4^ cryptococci in sterile saline was injected into the brain (e.g., bregma, cortex, striatum, or ventricle; stereotaxic coordinates are shown in Table 1) using a Hamilton syringe connected to a pump (model: UltraMicroPump3; World Precision Instruments). The skin incision on the dorsal head was closed with sterile nylon suture and 2-4% topical chlorhexidine solution was be applied over the closed incision. After the surgery, mice were placed on a clean recovery cage and monitored for survivability. The survival end points were inactivity, tachypnea, or loss of ≥ 25% of body weight from baseline weight. We monitored the mice twice daily for clinical signs, dehydration, and weight loss (Table 2). Animals showing signs of dehydration or that lost more than 10% weight received supportive care such as 1 mL of parenteral fluid supplementation (saline) and moist chow on the cage floor was provided. In separate infections, brain tissues were excised for processing for determination of CFU numbers and histopathological studies.

### Photographs

The surgical procedure, i.c. injections, physical changes (e.g., hydrocephalus), and excised brains were photographed with a digital camera and the pictures were transferred to a secured computer. We followed the recommendations from the University of Florida Animal Care Services Policy.

### CFU determinations

Brains were excised from euthanized mice and weighed 3- and 7-dpi. The brain tissue was homogenized in 5 mL of sterile phosphate buffered saline, serially diluted, a 100 μL suspension was plated on Sabouraud dextrose agar (BD Difco) and incubated at 30°C for 48 h. Quantification of viable yeast cells from infected animals were determined by CFU counting.

### Histological examinations

Brain tissue was excised from euthanized mice 7- and 20-dpi and fixed in 4% paraformaldehyde (Sigma) for 24 h. Tissues were managed, embedded in paraffin, and 4 μm coronal sections were cut and fixed to glass slides and then subjected either to hematoxylin and eosin (H&E) stain, to PAS stain, or to Prussian blue stain to examine host tissue, fungal morphology, or hemosiderosis, respectively. The tissues were then stained for GXM, microglia, and astrocytes using a mAb 18B7 (dilution: 1:1,000; anti-cryptococcal GXM IgG_1_ generated and generously provided by Arturo Casadevall at Johns Hopkins Bloomberg School of Public Health), Iba-1 (dilution: 1:1,000; FujiFilm Wako Chemicals USA Corp.), and GFAP (dilution: 1:1,000; DAKO), respectively, conjugated to horseradish peroxidase. For histopathological evaluations, whole slides scanning was performed using the Leica Aperio Scanscope CS and analyzed using The ImageScope software (version 12.4.3). Moreover, each slide was carefully examined with a Leica DMi8 inverted microscope, and photographed with a Leica DFC7000 T digital camera using the Leica software platform LAS X. Each image was blindly analyzed by three independent investigators. The pathological similarities observed in mice and humans with cerebral cryptococcosis were reported (Table 3) by a veterinary pathologist (Mohamed F. Hamed) and physician specialized in infectious diseases (Mircea R. Mihu) who independently analyzed the data.

**Table 3.**
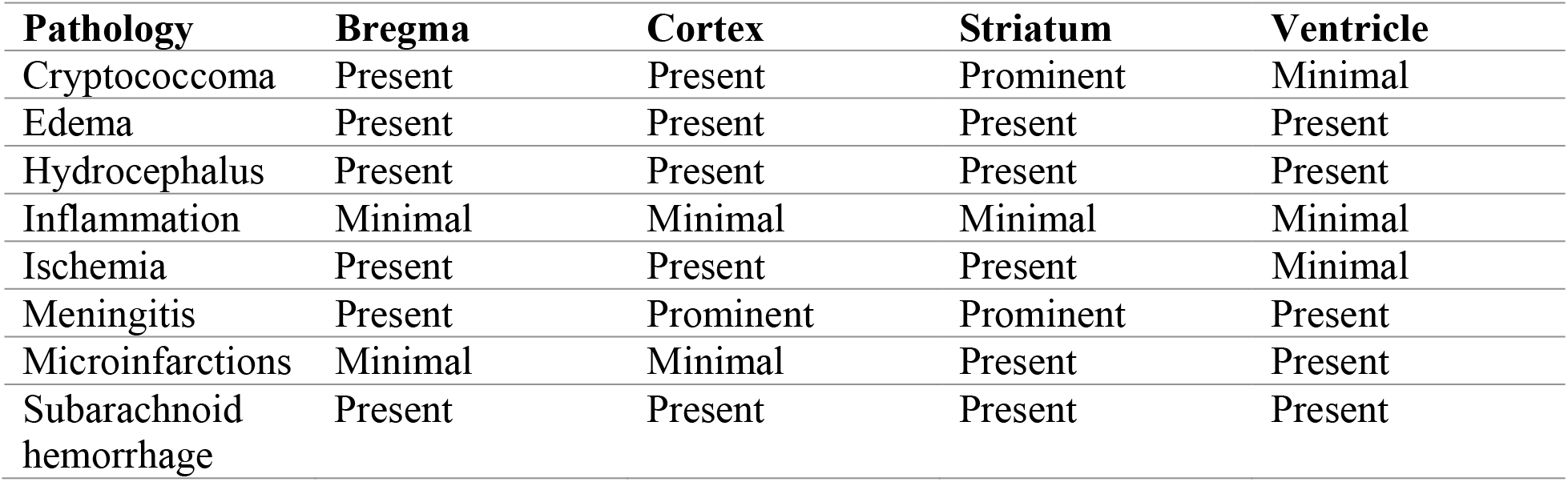
Pathological similarities observed in mice and humans with cerebral cryptococcosis.

| **Pathology** | **Bregma** | **Cortex** | **Striatum** | **Ventricle** |
| --- | --- | --- | --- | --- |
| Cryptococcoma | Present | Present | Prominent | Minimal |
| Edema | Present | Present | Present | Present |
| Hydrocephalus | Present | Present | Present | Present |
| Inflammation | Minimal | Minimal | Minimal | Minimal |
| Ischemia | Present | Present | Present | Minimal |
| Meningitis | Present | Prominent | Prominent | Present |
| Microinfarctions | Minimal | Minimal | Present | Present |
| Subarachnoid hemorrhage | Present | Present | Present | Present |

### Analysis of GXM and glia colocalization in brain tissue

Images of the brain tissue stained with GXM-, Iba-1 (microglia)-, and GFAP (astrocytes)- specific mAbs were analyzed to correlate GXM to glia colocalization by comparing differences in stain intensity in select regions of mice infected with *C. neoformans* in bregma, cortex, striatum, and ventricle using the NIH ImageJ software (version 1.46r). Each image was sectioned by a 6 × 8 mm square grid, and regions of each tissue sample were selected for analysis of unweighted light intensity using the ImageJ histogram function. The mean light intensities and standard deviations were recorded to identify and compare regions of varying stain intensity between GXM and the microglia or astrocyte stains. Following acquisition of light intensity data for each corresponding section of the stained samples, the data was plotted in Prism 9.4 (GraphPad software). The results were then compared and tested for statistical significance using correlation analysis and simple linear regression.

### Statistical analysis

All data were subjected to statistical analysis using Prism 9.4 (GraphPad). Differences in survival rates were analyzed by the log-rank test (Mantel-Cox). *P* values for multiple comparisons were calculated by analysis of variance (ANOVA) and were adjusted by use of the Tukey’s post-hoc analysis. *P* values for individual comparisons were calculated using student’s or multiple *t*-test analyses. Concentration-response curve for vascular reactivity was log-transformed, normalized to percent maximal response, and fitted using a nonlinear regression. *P* values of <0.05 were considered significant.

## SUPPLEMENTAL FIGURES

**Supplemental Fig. 1.**
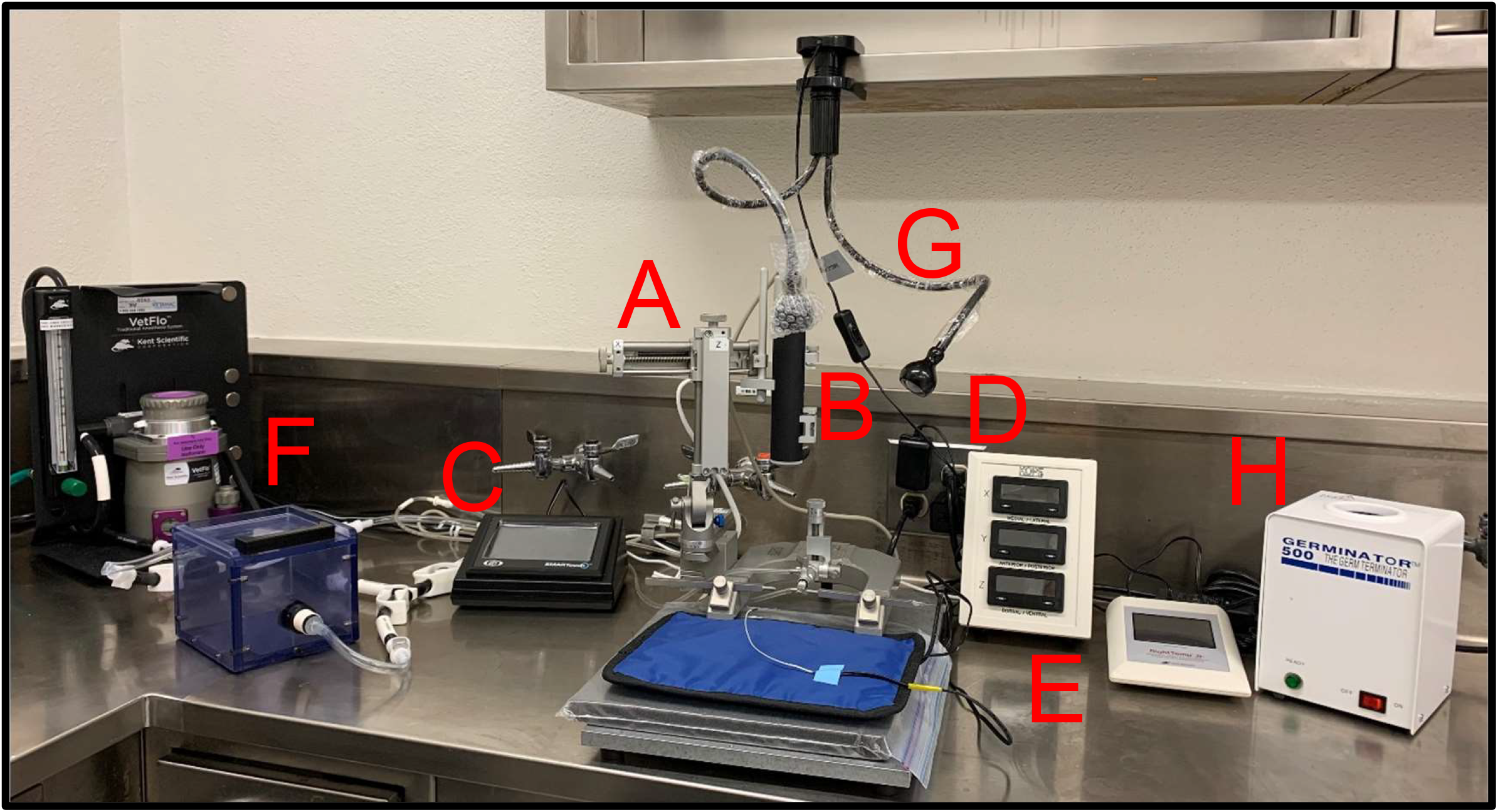
Stereotaxic apparatus and equipment setup used for i.c. *C. neoformans* infections in different regions of the brain of C57BL/6 mice. **(A)** Stereotaxic instrument uses a three-dimensional coordinate system (e.g., The Allen Mouse Brain Atlas; https://mouse.brain-map.org/static/atlas) to locate specific targets inside the mouse brain. **(B)** Ultra micro pump holds microinjection syringes to deliver picoliter to milliliter volumes and mounts directly on the stereotaxic frame. **(C)** Touch screen controller dispenses specific inoculum volume in a microinjection syringe into determined region of the mouse brain. **(D)** Digital display console design ensures full range of motion, angulation, and rotation of the stereotaxic manipulator. **(E)** Infrared warming pad with touchscreen controller and temperature feedback accurately monitors the mouse’s temperature. **(F)** Isoflurane anesthesia machine and chamber. **(G)** LED clamp light lamp with flexible gooseneck. **(H)** The stainless-steel glass bead bath decontaminates micro-dissecting instruments between procedures.

## ACKNOWLEDGEMENTS

M.F.H., M.E.M., C.L.C-N and L.R.M. were supported by the National Institute of Allergy and Infectious Diseases (NIAID award # R01AI145559) of the US National Institutes of Health (NIH). V.E. was supported by the UFCD’s Comprehensive Training Program in Oral Biology (NIH NIDCR Award # T90DE021990/R90DE022530). The funders had no role in the study design, data collection and analysis, decision to publish, or preparation of the manuscript.

## AUTHORSHIP CONTRIBUTIONS

All authors contributed to the project design and experimental procedures, analyzed data, provided the figure presentation, and manuscript writing.

## REFERENCES

1. Rajasingham R, Smith RM, Park BJ, Jarvis JN, Govender NP, Chiller TM, Denning DW, Loyse A, Boulware DR. 2017. Global burden of disease of HIV-associated cryptococcal meningitis: an updated analysis. Lancet Infect Dis 17:873–881.

2. Kontoyiannis DP, Marr KA, Park BJ, Alexander BD, Anaissie EJ, Walsh TJ, Ito J, Andes DR, Baddley JW, Brown JM, Brumble LM, Freifeld AG, Hadley S, Herwaldt LA, Kauffman CA, Knapp K, Lyon GM, Morrison VA, Papanicolaou G, Patterson TF, Perl TM, Schuster MG, Walker R, Wannemuehler KA, Wingard JR, Chiller TM, Pappas PG. 2010. Prospective surveillance for invasive fungal infections in hematopoietic stem cell transplant recipients, 2001-2006: overview of the Transplant-Associated Infection Surveillance Network (TRANSNET) Database. Clin Infect Dis 50:1091–100.

3. Sun HY, Wagener MM, Singh N. 2009. Cryptococcosis in solid-organ, hematopoietic stem cell, and tissue transplant recipients: evidence-based evolving trends. Clin Infect Dis 48:1566–76.

4. Levitz SM. 1991. The ecology of Cryptococcus neoformans and the epidemiology of cryptococcosis. Rev Infect Dis 13:1163–9.

5. Neilson JB, Fromtling RA, Bulmer GS. 1977. Cryptococcus neoformans: size range of infectious particles from aerosolized soil. Infect Immun 17:634–8.

6. Husain S, Wagener MM, Singh N. 2001. Cryptococcus neoformans infection in organ transplant recipients: variables influencing clinical characteristics and outcome. Emerg Infect Dis 7:375–81.

7. Singh N, Alexander BD, Lortholary O, Dromer F, Gupta KL, John GT, del Busto R, Klintmalm GB, Somani J, Lyon GM, Pursell K, Stosor V, Munoz P, Limaye AP, Kalil AC, Pruett TL, Garcia-Diaz J, Humar A, Houston S, House AA, Wray D, Orloff S, Dowdy LA, Fisher RA, Heitman J, Wagener MM, Husain S, Cryptococcal Collaborative Transplant Study G. 2007. Cryptococcus neoformans in organ transplant recipients: impact of calcineurin-inhibitor agents on mortality. J Infect Dis 195:756–64.

8. Dromer F, Mathoulin-Pelissier S, Launay O, Lortholary O, French Cryptococcosis Study G. 2007. Determinants of disease presentation and outcome during cryptococcosis: the CryptoA/D study. PLoS Med 4:e21.

9. Lortholary O, Improvisi L, Rayhane N, Gray F, Fitting C, Cavaillon JM, Dromer F. 1999. Cytokine profiles of AIDS patients are similar to those of mice with disseminated Cryptococcus neoformans infection. Infect Immun 67:6314–20.

10. Chretien F, Lortholary O, Kansau I, Neuville S, Gray F, Dromer F. 2002. Pathogenesis of cerebral Cryptococcus neoformans infection after fungemia. J Infect Dis 186:522–30.

11. Chang YC, Stins MF, McCaffery MJ, Miller GF, Pare DR, Dam T, Paul-Satyaseela M, Kim KS, Kwon-Chung KJ. 2004. Cryptococcal yeast cells invade the central nervous system via transcellular penetration of the blood-brain barrier. Infect Immun 72:4985–95.

12. Charlier C, Nielsen K, Daou S, Brigitte M, Chretien F, Dromer F. 2009. Evidence of a role for monocytes in dissemination and brain invasion by Cryptococcus neoformans. Infect Immun 77:120–7.

13. Eugenin EA, Greco JM, Frases S, Nosanchuk JD, Martinez LR. 2013. Methamphetamine alters blood brain barrier protein expression in mice, facilitating central nervous system infection by neurotropic Cryptococcus neoformans. J Infect Dis 208:699–704.

14. Santiago-Tirado FH, Onken MD, Cooper JA, Klein RS, Doering TL. 2017. Trojan Horse Transit Contributes to Blood-Brain Barrier Crossing of a Eukaryotic Pathogen. mBio 8.

15. Buchanan KL, Murphy JW. 1998. What makes Cryptococcus neoformans a pathogen? Emerg Infect Dis 4:71–83.

16. Fromtling RA, Shadomy HJ, Jacobson ES. 1982. Decreased virulence in stable, acapsular mutants of cryptococcus neoformans. Mycopathologia 79:23–9.

17. Littman ML. 1958. Capsule synthesis by Cryptococcus neoformans. Trans N Y Acad Sci 20:623–48.

18. Littman ML, Tsubura E. 1959. Effect of degree of encapsulation upon virulence of Cryptococcus neoformans. Proc Soc Exp Biol Med 101:773–7.

19. Rivera J, Feldmesser M, Cammer M, Casadevall A. 1998. Organ-dependent variation of capsule thickness in Cryptococcus neoformans during experimental murine infection. Infect Immun 66:5027–30.

20. Granger DL, Perfect JR, Durack DT. 1985. Virulence of Cryptococcus neoformans. Regulation of capsule synthesis by carbon dioxide. J Clin Invest 76:508–16.

21. Zaragoza O, Fries BC, Casadevall A. 2003. Induction of capsule growth in Cryptococcus neoformans by mammalian serum and CO(2). Infect Immun 71:6155–64.

22. Dykstra MA, Friedman L, Murphy JW. 1977. Capsule size of Cryptococcus neoformans: control and relationship to virulence. Infect Immun 16:129–35.

23. Jacobson ES, Tingler MJ, Quynn PL. 1989. Effect of hypertonic solutes upon the polysaccharide capsule in Cryptococcus neoformans. Mycoses 32:14–23.

24. Vartivarian SE, Anaissie EJ, Cowart RE, Sprigg HA, Tingler MJ, Jacobson ES. 1993. Regulation of cryptococcal capsular polysaccharide by iron. J Infect Dis 167:186–90.

25. Cherniak R, Sundstrom JB. 1994. Polysaccharide antigens of the capsule of Cryptococcus neoformans. Infect Immun 62:1507–12.

26. Goldman DL, Lee SC, Casadevall A. 1995. Tissue localization of Cryptococcus neoformans glucuronoxylomannan in the presence and absence of specific antibody. Infect Immun 63:3448–53.

27. Vecchiarelli A. 2000. Immunoregulation by capsular components of Cryptococcus neoformans. Med Mycol 38:407–17.

28. Pettoello-Mantovani M, Casadevall A, Smarnworawong P, Goldstein H. 1994. Enhancement of HIV type 1 infectivity in vitro by capsular polysaccharide of Cryptococcus neoformans and Haemophilus influenzae. AIDS Res Hum Retroviruses 10:1079–87.

29. Bicanic T, Harrison TS. 2004. Cryptococcal meningitis. Br Med Bull 72:99–118.

30. Kambugu A, Meya DB, Rhein J, O’Brien M, Janoff EN, Ronald AR, Kamya MR, Mayanja-Kizza H, Sande MA, Bohjanen PR, Boulware DR. 2008. Outcomes of cryptococcal meningitis in Uganda before and after the availability of highly active antiretroviral therapy. Clin Infect Dis 46:1694–701.

31. Nussbaum JC, Jackson A, Namarika D, Phulusa J, Kenala J, Kanyemba C, Jarvis JN, Jaffar S, Hosseinipour MC, Kamwendo D, van der Horst CM, Harrison TS. 2010. Combination flucytosine and high-dose fluconazole compared with fluconazole monotherapy for the treatment of cryptococcal meningitis: a randomized trial in Malawi. Clin Infect Dis 50:338–44.

32. Lee SC, Dickson DW, Casadevall A. 1996. Pathology of cryptococcal meningoencephalitis: analysis of 27 patients with pathogenetic implications. Hum Pathol 27:839–47.

33. Loyse A, Wainwright H, Jarvis JN, Bicanic T, Rebe K, Meintjes G, Harrison TS. 2010. Histopathology of the arachnoid granulations and brain in HIV-associated cryptococcal meningitis: correlation with cerebrospinal fluid pressure. AIDS 24:405–10.

34. Bicanic T, Brouwer AE, Meintjes G, Rebe K, Limmathurotsakul D, Chierakul W, Teparrakkul P, Loyse A, White NJ, Wood R, Jaffar S, Harrison T. 2009. Relationship of cerebrospinal fluid pressure, fungal burden and outcome in patients with cryptococcal meningitis undergoing serial lumbar punctures. AIDS 23:701–6.

35. Freitas I, Salazar T, Rodrigues P, Vilela M, Duarte A. 2022. An Uncommon Presentation of Cryptococcal Meningoencephalitis. Cureus 14:e21984.

36. Kouame-Assouan AE, Cowppli-Bony P, Aka-Anghui Diarra E, Assi B, Doumbia M, Diallo L, Adjien KC, Akani E, Sonan T, Diagana M, Boa YE, Kouassi B. 2007. [Two cases of cryptococcal meningitis revealed by an ischemic stroke]. Bull Soc Pathol Exot 100:15–6.

37. Sico JJ, Hughes E. 2006. Necrotising cryptococcal vasculitis in an HIV-negative woman. Mycoses 49:152–4.

38. Klock C, Cerski M, Goldani LZ. 2009. Histopathological aspects of neurocryptococcosis in HIV-infected patients: autopsy report of 45 patients. Int J Surg Pathol 17:444–8.

39. Jain KK, Mittal SK, Kumar S, Gupta RK. 2007. Imaging features of central nervous system fungal infections. Neurol India 55:241–50.

40. Lee SC, Dickson DW, Brosnan CF, Casadevall A. 1994. Human astrocytes inhibit Cryptococcus neoformans growth by a nitric oxide-mediated mechanism. J Exp Med 180:365–9.

41. Denning DW, Armstrong RW, Lewis BH, Stevens DA. 1991. Elevated cerebrospinal fluid pressures in patients with cryptococcal meningitis and acquired immunodeficiency syndrome. Am J Med 91:267–72.

42. Fries BC, Lee SC, Kennan R, Zhao W, Casadevall A, Goldman DL. 2005. Phenotypic switching of Cryptococcus neoformans can produce variants that elicit increased intracranial pressure in a rat model of cryptococcal meningoencephalitis. Infect Immun 73:1779–87.

43. Mishra AK, Arvind VH, Muliyil D, Kuriakose CK, George AA, Karuppusami R, Benton Carey RA, Mani S, Hansdak SG. 2018. Cerebrovascular injury in cryptococcal meningitis. Int J Stroke 13:57–65.

44. Gibson JF, Bojarczuk A, Evans RJ, Kamuyango AA, Hotham R, Lagendijk AK, Hogan BM, Ingham PW, Renshaw SA, Johnston SA. 2022. Blood vessel occlusion by Cryptococcus neoformans is a mechanism for haemorrhagic dissemination of infection. PLoS Pathog 18:e1010389.

45. Blasi E, Barluzzi R, Mazzolla R, Bistoni F. 1993. Differential host susceptibility to intracerebral infections with Candida albicans and Cryptococcus neoformans. Infect Immun 61:3476–81.

46. Blasi E, Barluzzi R, Mazzolla R, Mosci P, Bistoni F. 1992. Experimental model of intracerebral infection with Cryptococcus neoformans: roles of phagocytes and opsonization. Infect Immun 60:3682–8.

47. Blasi E, Mazzolla R, Barluzzi R, Mosci P, Bistoni F. 1994. Anticryptococcal resistance in the mouse brain: beneficial effects of local administration of heat-inactivated yeast cells. Infect Immun 62:3189–96.

48. Mazzolla R, Barluzzi R, Brozzetti A, Boelaert JR, Luna T, Saleppico S, Bistoni F, Blasi E. 1997. Enhanced resistance to Cryptococcus neoformans infection induced by chloroquine in a murine model of meningoencephalitis. Antimicrob Agents Chemother 41:802–7.

49. Kinnecom K, Pachter JS. 2005. Selective capture of endothelial and perivascular cells from brain microvessels using laser capture microdissection. Brain Res Brain Res Protoc 16:1–9.

50. Olszewski MA, Noverr MC, Chen GH, Toews GB, Cox GM, Perfect JR, Huffnagle GB. 2004. Urease expression by Cryptococcus neoformans promotes microvascular sequestration, thereby enhancing central nervous system invasion. Am J Pathol 164:1761–71.

51. Pollock H, Hutchings M, Weller RO, Zhang ET. 1997. Perivascular spaces in the basal ganglia of the human brain: their relationship to lacunes. J Anat 191 (Pt 3):337–46.

52. Lee SC, Casadevall A. 1996. Polysaccharide antigen in brain tissue of AIDS patients with cryptococcal meningitis. Clin Infect Dis 23:194–5.

53. Lee SC, Casadevall A, Dickson DW. 1996. Immunohistochemical localization of capsular polysaccharide antigen in the central nervous system cells in cryptococcal meningoencephalitis. Am J Pathol 148:1267–74.

54. Barluzzi R, Brozzetti A, Delfino D, Bistoni F, Blasi E. 1998. Role of the capsule in microglial cell-Cryptococcus neoformans interaction: impairment of antifungal activity but not of secretory functions. Med Mycol 36:189–97.

55. Lipovsky MM, Gekker G, Hu S, Ehrlich LC, Hoepelman AI, Peterson PK. 1998. Cryptococcal glucuronoxylomannan induces interleukin (IL)-8 production by human microglia but inhibits neutrophil migration toward IL-8. J Infect Dis 177:260–3.

56. Ellerbroek PM, Hoepelman AI, Wolbers F, Zwaginga JJ, Coenjaerts FE. 2002. Cryptococcal glucuronoxylomannan inhibits adhesion of neutrophils to stimulated endothelium in vitro by affecting both neutrophils and endothelial cells. Infect Immun 70:4762–71.

57. Ellerbroek PM, Ulfman LH, Hoepelman AI, Coenjaerts FE. 2004. Cryptococcal glucuronoxylomannan interferes with neutrophil rolling on the endothelium. Cell Microbiol 6:581–92.

58. Ellis JP, Kalata N, Joekes EC, Kampondeni S, Benjamin LA, Harrison TS, Lalloo DG, Heyderman RS. 2018. Ischemic stroke as a complication of cryptococcal meningitis and immune reconstitution inflammatory syndrome: a case report. BMC Infect Dis 18:520.

59. Shimoda Y, Ohtomo S, Arai H, Ohtoh T, Tominaga T. 2017. Subarachnoid small vein occlusion due to inflammatory fibrosis-a possible mechanism for cerebellar infarction in cryptococcal meningoencephalitis: a case report. BMC Neurol 17:157.

60. Chen SHM, Stins MF, Huang SH, Chen YH, Kwon-Chung KJ, Chang Y, Kim KS, Suzuki K, Jong AY. 2003. Cryptococcus neoformans induces alterations in the cytoskeleton of human brain microvascular endothelial cells. J Med Microbiol 52:961–970.

61. Easterly ME, Foltz CJ, Paulus MJ. 2001. Body condition scoring: comparing newly trained scorers and micro-computed tomography imaging. Lab Anim (NY) 30:46–9.

